# Fast and accurate local ancestry inference with Recomb-Mix

**DOI:** 10.1101/2023.11.17.567650

**Authors:** Yuan Wei, Degui Zhi, Shaojie Zhang

## Abstract

The availability of large genotyped cohorts brings new opportunities for revealing the high-resolution genetic structure of admixed populations via local ancestry inference (LAI), the process of identifying the ancestry of each segment of an individual haplotype. Though current methods achieve high accuracy in standard cases, LAI is still challenging when reference populations are more similar (e.g., intra-continental), when the number of reference populations is too numerous, or when the admixture events are deep in time, all of which are increasingly unavoidable in large biobanks. Here, we present a new LAI method, Recomb-Mix. Recomb-Mix integrates the elements of existing methods of the site-based Li and Stephens model and introduces a new graph collapsing trick to simplify counting paths with the same ancestry label readout. Through comprehensive benchmarking on various simulated datasets, we show that Recomb-Mix is more accurate than existing methods in diverse sets of scenarios while being competitive in terms of resource efficiency. We expect that Recomb-Mix will be a useful method for advancing genetics studies of admixed populations.

## Introduction

Local ancestry inference (LAI) is a process of assigning the ancestral population labels of each segment on an individual’s genome sequence. LAI is not only useful for better study of human demographic history (Martin et al. 2017) but also can enable several downstream tasks, including admixture mapping (Reich et al. 2005), ancestry-aware genome-wide association studies (GWAS) (Pasaniuc et al. 2011), and ancestry-specific polygenic risk scores (Duncan et al. 2019). Recent studies show that local ancestry information improves the resolution of association signals in GWAS (Atkinson et al. 2021), helping to infer the high-resolution of genomic regions containing genes as under selection (Hamid et al. 2023). Local ancestry calls contribute to understanding the impact of genetic variants that cause disease (Hou et al. 2023) and the accuracy of polygenic scores of genetically based predictions (Ding et al. 2023).

The recent availability of biobank-scale genotyped datasets (Bycroft et al. 2018; Kurki et al. 2023) and the rising of enormous databases from direct-to-consumer genetic companies (Durand et al. 2021; Wang et al. 2021) create new challenges and opportunities for LAI. Participants in biobanks may be from highly imbalanced source populations. Inferring ancestral components underrepresented in these reference populations is in great need. On the other hand, the admixture to be inferred in the samples may be multi-ways and may be from both recent and distant admixture events. However, opportunities coexist with such challenges. More diverse samples, e.g., the Human Genome Diversity Project (HGDP) (Bergström et al. 2020), are becoming available as reference panels. The number of participants in biobanks is much larger and more representative than in previous reference panels, and thus, more potential sub-continental ancestral information becomes available from biobanks. Although methods for revealing sub-continental or even sub-population clusters are available (e.g., (Lawson et al. 2012)), they are mostly non-LAI methods and only capture global ancestry. With the availability of diverse samples in biobanks and the need for in-depth knowledge of admixed individuals, current LAI methods are unable to capitalize on these opportunities fully.

Existing LAI methods fall into two categories: site-based and window-based. Originally, Hap-Mix (Price et al. 2009), based on the extended Li and Stephens (LS) Hidden Markov model (Li and Stephens 2003), is developed to model different transition probabilities for within population and between population jumps. To make it tolerate mismatches, emission probabilities are also introduced, and the model is not very efficient and cannot be scaled up (Geza et al. 2018). Later, window-based methods are gaining popularity (e.g., RFMix (Maples et al. 2013), G-Nomix (Hilmarsson et al. 2021), SALAI-Net (Oriol Sabat et al. 2022)). These methods take short stretches of sites as windows and define window-based features. It first makes local ancestry prediction over each window and then uses certain post-processing to smooth out the labels across all windows. For each window, a certain machine-learning approach is typically used. However, the predefined window boundaries are not necessarily optimized, and the noisy initial window labels can be difficult to correct by post-processing. Loter (Dias-Alves et al. 2018) is a recent site-based method under the LS framework. It formulates the LAI problem as a combinatorial best-path problem in a graph, which can be solved efficiently by dynamic programming. However, its problem formulation is simplistic in that it does not take into account the useful information encoded in the LS model, such as differential transition probabilities for within and between populations and variable recombination rates across sites. Loter reported that it underperformed RFMix (Maples et al. 2013) and LAMP-LD (Baran et al. 2012) for datasets with recent admixture events (i.e. *<* 150 generations). Therefore, there is room for improvement over the Loter approach by introducing an LS-inspired parametrization of its scoring function.

In this study, we developed Recomb-Mix, a site-based method that is both accurate and efficient.

Our main insight is that we do not have to have an exact LS formulation. The gist of the HapMix LS model is the differential transition penalties for within and between populations and assigning the population labels for a site by comparing the paths going through it versus the paths by-passing it. We achieve the same spirit by setting the within-population transition penalty to zero and collapsing the nodes representing the allele values at each site. This allows both run time and space efficiency while achieving higher accuracy than Loter. Recomb-Mix is designed to have robustness, scalability, and superior accuracy on LAI. Applications to real human datasets confirmed the genetic differences among populations and provided potential explanations for how the evolutionary processes shape these differences.

## Methods

### The Li and Stephens framework for site-based local ancestry inference

Recomb-Mix is inspired by HapMix (Price et al. 2009) and Loter (Dias-Alves et al. 2018), all of which are under the Li and Stephens (LS) framework. These two methods extend the basic LS model (Li and Stephens 2003) to capture the difference between inter-population and intra-population transition probabilities. The LS model defines the conditional probability of any haplotype sequence given a set of haplotypes in a panel as a Hidden Markov Model (HMM). The states of the HMM are individual sequence positions in the panel. By treating all haplotypes equally, transition probability in the HMM can be specified by just the probability of switching a haplotype template or staying at the same template. In a non-probabilistic combinatorial formulation, transition probability can be modeled as a template change penalty. HapMix extends the model to have population labels as augmented HMM state labels and introduces two transition probability parameters, i.e., small-scale (between haplotypes from within a reference population) and large-scale (between the reference populations). See equations (0.1) and (0.2) in the HapMix paper for the detailed definition of the population-label-aware LS model. Loter formulates LAI as a graph optimization problem that finds a best-scoring path over a site-level graph. It can be viewed as an LS “copying model” with simplified non-probabilistic parameterization. Loter applies the same penalty to haplotype template switches in both cases within or across reference populations.

Through the unified view of the LS framework and the graph optimization formulation (Table 1), Recomb-Mix introduces special parameterization to the LS model to induce graph simplification and more biologically relevant scoring function. First, by assuming no template change penalty when switching haplotype templates within a reference population, Recomb-Mix enables the collapsing of the reference panel to a compact population graph. Generating a compact population graph greatly reduces the size of reference populations and retains the ancestry information, as genetic markers having the same allele values per population are collapsed in the compact population graph. Different template change penalties are used when switching haplotype templates within a reference population and between the reference populations. The template change penalty within a reference population is set to zero, and recombination rates from a genetic map parameterize the template change penalty between the reference populations. Second, Recomb-Mix’s scoring function Σ(*d* + *w* · *r*) is a simplified version of HapMix’s. Still, it is a richer version than that of Loter’s. In Recomb-Mix’s scoring function, *d* is a mismatch penalty score regarding a site on the query haplotype and the corresponding site in the reference panel, *w* is the weight for the relative importance of the mismatch cost and the recombination cost, and *r* is a normalized recombination rate penalty score between two sites translated from the genetic distance. Table 1 shows the differences in scoring functions between some LAI methods under the LS framework. HapMix incorporates recombination, miscopying, mutation, and genetic distance parameters into its scoring function. With such a number of required biological parameters, lacking accurate population information may lead to biased inference results; that is, the estimated biological parameters required for HapMix may not be the correct parameters for the given dataset, and the ancestry inference result based on such parameters is influenced (Patin et al. 2014; Suarez-Pajes et al. 2021). In contrast to HapMix’s complex scoring function, Loter uses a simple one that does not adopt any recombination information. Recomb-Mix takes Loter’s simplicity but adds the notion of genetic distance to encode genetic information, to achieve high computability and accuracy simultaneously.

**Table 1:** Comparison of scoring functions in HapMix (Price et al. 2009), Loter (Dias-Alves et al. 2018), and Recomb-Mix (this study) under the Li and Stephens (LS) framework.

| Scoring Function | HapMix | Loter | Recomb-Mix |
| --- | --- | --- | --- |
| Within population transition probability / template change penalty | Recombination parameter | 1 | 0 |
| Across population transition probability / template change penalty | Recombination parameter and miscopying parameter | 1 | $r$, recombination penalty |
| Emission probability / mismatch penalty | Mutation parameter | Mismatch penalty | $d$, mismatch penalty |
| Parameterization of transition probability | $1 - e^p$, where $p$ is the product of genetic distance and recombination parameter | Bootstrap aggregation (bagging) | $w$, weight |

Besides the scoring function, Recomb-Mix has other differences from HapMix and Loter. Recomb-Mix and Loter can handle multi-way admixture inference, as HapMix can only tackle two-way admixture. Both Recomb-Mix and HapMix use genetic map (Church et al. 2011; Schneider et al. 2017) to help out the inference, while Loter does not take any biological information as input. From the HMM algorithm perspective, HapMix uses the forward-backward algorithm to update transition and emission probabilities and then estimate the hidden ancestral states (Price et al. 2009; Wu et al. 2021). Loter’s approach minimizes an objective function using dynamic programming, a Viterbi-like algorithm (Dias-Alves et al. 2018; Oriol Sabat et al. 2022). Recomb-Mix takes advantage of the graph optimization formulation as Loter does but keeps population-level information only in a compact population graph. All possible paths in the graph can be viewed as a set of “combined” paths from the original graph that emit the same population label readout. Thus, it has a flavor of a “forward-like” algorithm as an ancestry label is assigned to an individual node according to “combined” paths passing through it.

### Recomb-Mix

The Recomb-Mix method is inspired by the LS HMM and implemented using a graph optimization approach. Like the LS model, it assumes that an admixed individual haplotype is modeled as a mosaic of individual haplotypes from a reference panel. Recomb-Mix constructs a population graph from a given reference panel to infer the ancestral label at each locus on a given admixed individual haplotype by finding a threading path that resembles the admixed individual haplotype the most among all the paths. In the population graph, the allele values of individual haplotypes are grouped by each site. Then, the population graph is transformed into a compact population graph by collapsing the site nodes with the same allele value and ancestral label into one node. A mismatch penalty at each site occurs when there is a difference between the collapsed site nodes’ allele value and the corresponding site’s allele value in the admixed individual haplotype. The collapsed site nodes are linked to population emission nodes based on the ancestral label of each node, and the population emission nodes make a cross-population connection to the population emission nodes of the next site. A template change penalty regarding recombinations between each site occurs when two population emission nodes of adjacent sites having different ancestral labels are connected. Then, the population emission nodes are expanded to genotype emission nodes linked to the site nodes for the next site. This process is similar to a “forward-like” approach as each node in the compact population graph can be viewed as a bundle of nodes in the original population graph being consolidated as one, and their ancestral labels are assigned by all low penalty threading paths passing through it. Recomb-Mix sums over mismatch penalties and template change penalties through all possible threading paths to determine the one that has the minimum penalty score. The ancestral label of each site in the admixed individual haplotype is assigned the same ancestral label of each corresponding node on such threading path.

To formally define the Recomb-Mix method, a reference panel having *m* individual haplotypes with *n* sites can be transformed as a population graph *G* = (*V, E*), representing the HMM of this set of haplotypes. *V* is the union of the starting and the ending nodes {*s, e*} and the site nodes *S_j_* for *j* ∈ [1*, n*]. 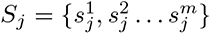 is the set of nodes representing alleles of haplotypes at position *j*. *E* is the union of the edges from 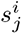 to 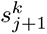 for all *j* ∈ [1*, n* − 1], *i, k* ∈ [1*, m*], and the edges from *s* to 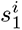 and 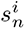 to *e* for *i* ∈ [1*, m*]. 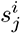 is the node that the *i*-th haplotype has at site *j*. Each node at every site of every haplotype has an associated ancestral label. It is assumed there are *p* populations presented in the reference panel, and each population has an ancestral label in [1*, p*]. The ancestral label of node 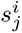 is 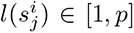. The allele value of node 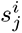 is 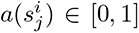, assuming all sites are bi-allelic. The population graph *G* is further transformed into a compact population graph *G′* = (*V ′, E′*) by collapsing all nodes with the same allele value and ancestral label to one node in every site. *V ′* is the union of the starting and the ending nodes {*s, e*} and the site nodes 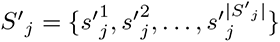 for *j* ∈ [1*, n*]. 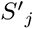 is a set of nodes representing all unique pairs of allele values and ancestral labels in *S_j_* (i.e., there is a node 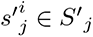 if and only if there is a node *s* ∈ *S_j_* such that 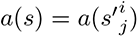 and 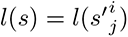 and for all 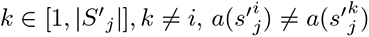 or 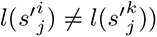. *E′* is the union of the edges from *u*_1_ to *u*_2_ for all 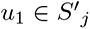 and 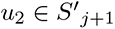 for *j* ∈ [1, *n* − 1], and the edges from *s* to *u*_1_ and *u*_2_ to *e* for all 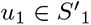 and 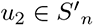.

To calculate all possible threading paths in *G′* for a query admixed individual haplotype *Q* with *n* sites, *Q* = (*q*_1_*, q*_2_*, … q_n_*) (the allele value of *q_i_* is *a*(*q_i_*) ∈ [0, 1]), Recomb-Mix incorporates the mismatch penalty and the template change penalty into its objective function. The mismatch penalty function is defined as *d*(*x*_1_*, x*_2_), where *x*_1_ and *x*_2_ are allele values. The template change penalty function is defined as *r*(*y*_1_*, y*_2_), where *y*_1_ and *y*_2_ are ancestral label values. Then, the cost of a candidate threading path *P* = (*u*_1_*, u*_2_*, …, u_n_*) is defined as:

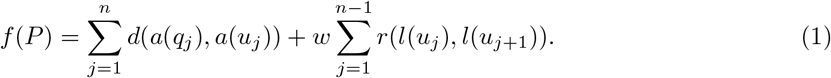

In Equation 1, *w* is a scale factor to balance the mismatch and recombination costs. Let *P ^∗^* be the threading path having the minimum penalty cost among all candidate threading paths in *G′*. The ancestral labels of the nodes in *P ^∗^* from *s* to *e* are the estimated ancestral labels of sites in *Q*, that is (*l*(*u^∗^*_1_)*, l*(*u^∗^*_2_)*, …, l*(*u^∗^_n_*)). Thus, LAI can be formulated as a problem to find *P ^∗^* in *G′*.

Figure 1 is an example of local ancestry inference with Recomb-Mix. A reference panel having seven individual haplotypes, eight sites, and two ancestral labels (shown in red and blue) is represented as a population graph *G*. Nodes representing each site are fully connected to nodes representing their adjacent site. A node *s* is connected to all nodes for the first site, and all nodes for the last site are connected to a node *e*. *Q* is a query of an admixed individual haplotype. *G* is then transformed into a compact population graph *G′*, and a threading path having the minimum penalty score is selected from node *s* to node *e* in *G′*, to be used to paint the admixed individual haplotype query by assigning the estimated ancestral label to each site in *Q*. Figure 1B is an example to show why *G′* is still an LS model. It demonstrates how nodes representing sites in positions three and four in *G* are transformed into the corresponding nodes in *G′*. First, the set of nodes representing alleles of haplotypes at position three (the first column) is freely collapsed to genotyping emission nodes (the second column) with the same allele value and ancestral label to one node. Second, the genotyping emission nodes are linked to population emission nodes (filled red and blue nodes in the third column) according to their ancestral labels. Then, the population emission nodes at position three make cross-population connections to the population emission nodes at position four (filled red and blue nodes in the fourth column). A penalty *r* is applied to the connections of population emission nodes when their ancestral labels differ. Finally, the population emission nodes at position four are linked to the set of nodes representing alleles of haplotypes at position four (the fifth column), and those nodes are freely collapsed to genotyping emission nodes (the sixth column).

**Figure 1:**
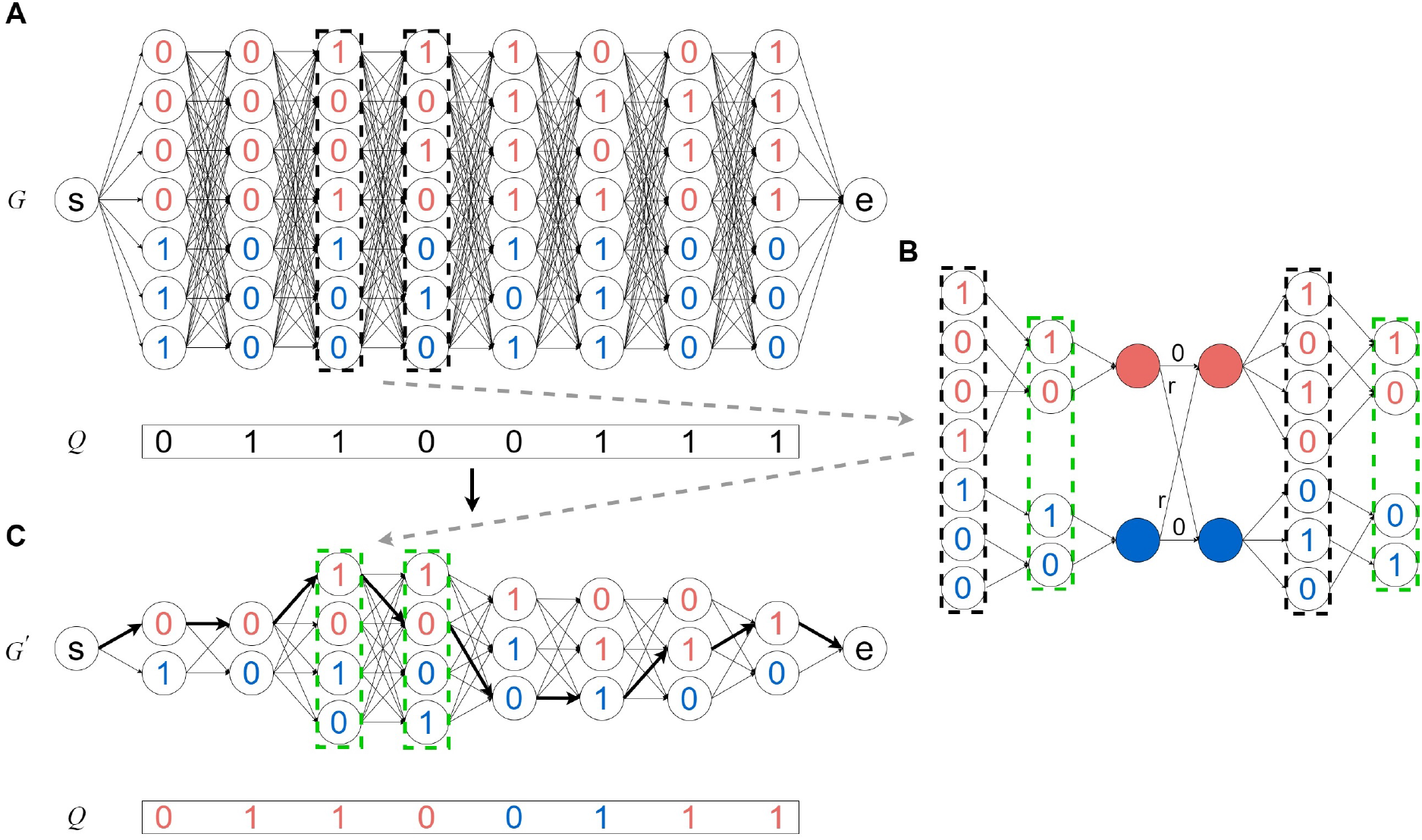
An example of local ancestry inference with Recomb-Mix. (A) *G* is a population graph representing the HMM in Recomb-Mix, constructed from a given reference panel. *G* contains seven haplotypes with eight sites belonging to two populations (shown in red and blue). *Q* is a query of an admixed individual haplotype. (B) A transformation process from nodes in sites three and four in G to nodes in the corresponding sites in *G′*. Nodes in black boxes correspond to the nodes in sites three and four in *G*. Nodes in green boxes correspond to the nodes in sites three and four in *G′*. The filled nodes in red and blue are population emission nodes in sites three and four. *r* is a cross-population penalty. (C) *G′* is a compact population graph transformed from *G*. *Q* is assigned with estimated ancestral labels for each site (shown in red and blue on allele values), according to a threading path selected with minimum penalty score (shown as bold edges) in *G′*.

Recomb-Mix uses a simplified scoring function (Equation 1) like the one HapMix uses to calculate the penalty score of each threading path. The mismatch penalty is determined by simply comparing the allele values of each site in *Q* to the corresponding nodes in *G′*. For each site, a mismatch penalty is applied if the allele values are not the same. *d*(·) in Equation 1 is implemented as *d*(*a*(*q_j_*)*, a*(*u_j_*)) = 0 if *a*(*q_j_*) = *a*(*u_j_*); otherwise *d*(*a*(*q_j_*)*, a*(*u_j_*)) = 1 for site *j* of *u_j_* in *G′* and *q_j_* in *Q*. The template change penalty is determined by the recombination rate between site *j* and *j* + 1 and the ancestral labels of *u_j_* and *u_j_*_+1_ in *G′*. Since the recombination rate between two sites is inversely proportional to the probability of the edge connecting these sites being a recombination breakpoint, the template change penalty cost is determined by the reciprocal of the recombination rate between the two sites. To leverage the linkage disequilibrium (LD) effect (haplotype information of allele correlations) from the LS model, the template change penalty is applied to the edge connecting two adjacent nodes if they have different ancestral labels. No template change penalty is applied if two adjacent nodes on a threading path share the same ancestral label. This setting allows the representation of a more diverse set of haplotypes than those explicitly listed in the panel. Not giving any penalties to them allows all possible threading paths jumping between templates within a population in *G* to be treated equally in *G′*. Of course, this setting is quite simplistic: it could risk allowing too much diversity. Also, it is possible to set the within-population template change penalty to a value other than zero, or some more sophisticated settings. However, setting this to zero captures the main idea of differentiating within-versus across-population transition probability. *r*(·) in Equation 1 is implemented as *r*(*l*(*u_j_*)*, l*(*u_j_*_+1_)) = 0 if *l*(*u_j_*) = *l*(*u_j_*_+1_); otherwise *r*(*l*(*u_j_*)*, l*(*u_j_*_+1_)) = *R_j,j_*_+1_ for site *j* and *j* +1 of *u_j_* and *u_j_*_+1_ in *G′*. *R_j,j_*_+1_ is the normalized reciprocal of the recombination rate between site *j* and *j* + 1. To calculate *R_j,j_*_+1_, min-max normalization is used to scale the range of the recombination rates from the genetic map into [0, 1]. The normalized recombination rates R*_j,j_*_+1_ are further processed by applying a reciprocal function to obtain *R_j,j_*_+1_, where *R_j,j_*_+1_ = 2*/*(R*_j,j_*_+1_ + 1). The range of the normalized reciprocal of the recombination rates is [1, 2]. The normalization step is to ensure the template change penalty is in the same order of magnitude as the mismatch penalty to prevent the domination of any penalties.

Representing the LS model as a compact population graph is efficient in terms of time and space. Recomb-Mix uses a dynamic programming approach to solve the problem of finding *P ^∗^* in *G′*. Using *G′* instead of *G* to compute the minimum penalty score among all possible threading paths significantly improves the computing time. The maximum indegree and outdegree of any node in *G′* is a constant value 2*p*, assuming all sites are bi-allelic and the number of populations presented in the reference panel is small (i.e., *p* ≪ *m*). For each site, each population’s minimum mismatch penalty score is tracked alongside the minimum mismatch penalty score over all the populations. The candidate threading paths of each node having the current minimum penalty score can be determined by comparing two candidate penalty scores of the adjacent site, which are the minimum penalty score whose population is the same and the minimum penalty score over all the populations. Thus, the time complexity of computing the penalty scores on *G′* is *O*(*np*). Using *G′* also substantially alleviates the demand for spaces to store a large reference panel. The space complexity is reduced from *O*(*nm*) to *O*(*np*), as *G′* stores at most 2*p* nodes per site. Using a compact population graph to reduce space usage is similar to but different from existing approaches in phasing and imputation. In a popular phasing method SHAPEIT (Delaneau et al. 2019), each node represents its allele value and each edge represents the weight of the number of individual haplotypes in its reference panel. This approach makes SHAPEIT have a space complexity of *O*(*nj*) (*j* is the number of conditioning states for each marker), which helps speed up its subsequent HMM calculation (Delaneau et al. 2012). Likewise, another popular phasing and imputation method, Beagle (Browning et al. 2018), constructs its HMM state space from its reference panel by leveraging composite reference haplotypes. The same haplotype compression technique is later adopted by FLARE (Browning et al. 2023). For each query individual haplotype, FLARE finds Identity-By-State (IBS) segments using Positional Burrows-Wheeler Transform (PBWT) algorithm (Durbin 2014) from the reference haplotypes and then makes composite reference haplotypes by stitching those IBS segments. Utilizing the composite reference haplotypes costs a *O*(*nm′*) space complexity, where *m′* depends on the number and locations of IBS segments of the query individual haplotype against the original reference panel. Usually, *m′* is relatively small, as it is expected there exist many long IBS segments.

Another advantage of presenting a reference panel as a compact population graph is that Recomb-Mix can process the reference panel regardless of whether the panel is phased or not. When converting a reference panel into a compact population graph, the order of the sites from two haplotypes of an individual is irrelevant, thanks to a property that the out-neighborhood of a node *u* in a graph is the set of nodes adjacent to *u*. Thus, Recomb-Mix is flexible to handle both phased and unphased reference panels.

### Simulated datasets

To evaluate the performance of Recomb-Mix, several admixture datasets were simulated using SLiM v4.0 (Haller and Messer 2019). The input data were individuals in Whole-Genome Sequencing (WGS) form from various populations in the study of the 1000 Genomes Project (TGP) (Auton et al. 2015; Clarke et al. 2016) and the Human Genome Diversity Project (HGDP) (Bergström et al. 2020). Each input population was split into disjoint sets of founders and references. The admixture population was simulated as the descendants of admixed founders from different populations. The admixed individuals were sampled from the admixture population, and the referenced individuals were sampled from each reference population. A set of three-way inter-continental datasets of Chromosome 18 were simulated using YRI, CEU, and CHB individuals (representing African, European, and Asian populations; more descriptions of the populations are available in Supplemental Table S1) from the TGP dataset. Various sizes of reference panels (i.e., 100, 250, 500, and 1,000) and numbers of generations after the admixture event (i.e., 15, 50, 100, and 200) were examined. The average recombination rate and mutation rate used for the simulation was 1.46455e-08 and 1.29e-08 per base pair per generation, according to the stdpopsim library (Adrion et al. 2020). A 0.02% genotyping error, following Browning et al. (2023), was added to admixed and reference individuals. The ground truth ancestral labels of admixed individuals were extracted from the SLiM output tree sequence (Haller et al. 2019). Additionally, a set of seven-way inter-continental datasets of Chromosome 18 were simulated using AFR, EAS, EUR, NAT, OCE, SAS, and WAS individuals (representing African, East Asian, European, American, Oceanian, Central and South Asian, and West Asian populations) from the HGDP dataset. The HGDP dataset was phased and imputed using Beagle 5.4 (Browning et al. 2018, 2021). The goal of simulating the seven-way admixture is to explore how well LAI methods are able to distinguish local ancestral segments from the admixture of a large number of ancestral populations.

To explore the power of LAI at the intra-continental level, a set of intra-continental datasets were simulated using TSI, FIN, and GBR individuals (representing Italian, Finnish, and British populations) using the same settings as the inter-continental datasets. To explore the influence of the uneven proportion of individuals per population in founders and references, two variations of the three-way 15-generation intra-continental datasets with uneven founders (i.e., 68 Italian, 32 Finnish, and 100 British individuals) or uneven references (i.e., 170 Italian, 80 Finnish, and 250 British individuals) were simulated. Both cases were 1/3, 1/6, and 1/2 individuals to the entire population panel. Additionally, an experiment was conducted to test the case when the reference panel size was ultra-small, i.e., the reference panel size was 20 and 50 (or only about 7 or 17 individuals per population in the reference panel).

### Benchmark setup

Two conventional measurements were used to evaluate the performance of LAI methods. The squared Pearson’s correlation coefficient *r*^2^ value (used by FLARE, LAMP-LD (Baran et al. 2012), and MOSAIC (Salter-Townshend and Myers 2019)) and the accuracy rate of the correctly-predicted markers (used by G-Nomix, Loter, RFMix, and SALAI-Net). The *r*^2^ value was followed by LAMP-LD’s definition (Baran et al. 2012), in which the *r*^2^ value is defined as the one between the true and the inferred number of alleles from each of the populations, averaged over all the populations. The criteria used by FLARE that markers were filtered with minor allele frequency ≤ 0.005 and minor allele count ≤ 50 (Browning et al. 2023) was also applied. *r*^2^ values are mainly reported in the benchmarks but accuracy rates are also available, mostly in the supplemental. It is found that the results of *r*^2^ values and accuracy rates are often consistent.

Recomb-Mix was tested against several datasets with the following LAI methods: FLARE (Browning et al. 2023), G-Nomix (Hilmarsson et al. 2021), Loter (Dias-Alves et al. 2018), RFMix (Maples et al. 2013), and SALAI-Net (Oriol Sabat et al. 2022). The weight parameter *w* was tuned to *w* = 1.5, which provides the best performance of Recomb-Mix for the given datasets. Parameters used for other methods are available in Supplemental Table S2. FLARE is the most recent proposed LAI method. It is a site-based generative method under the LS framework, extended from HapMix (Price et al. 2009), which models the hidden local ancestry of each site. FLARE shows encouraging speed and accuracy because its performance is optimized for computation resources, and its model is designed to be flexible for optional parameters to deal with various situations. G-Nomix is a window-based discriminative approach using two-layer prediction to perform LAI, in which it uses Logistic Regression as a base model and XGBoost as a smoother model. It is currently the leading LAI method due to its promising speed and accuracy. Loter frames the ancestry prediction problem as a graph optimization problem. It is prioritized for LAI on distant admixture events and good for non-model species as no biological information is required. RFMix, another window-based discriminative approach, is one of the popular LAI methods. It applies a conditional random field model to LAI, particularly using random forest classification. RFMix shows a robust performance on multi-way admixture datasets. SALAI-Net is also a window-based discriminative approach developed from its predecessor LAI-Net (Montserrat et al. 2020), the first neural network-based LAI. It uses a reference matching layer and a smoother layer (i.e., a combination of cosine similarity score and neural network) to perform LAI. In addition to the adoption of GPU hardware, SALAI-Net utilizes a pre-trained generalized model, making it free from re-training and parameter tuning during the inference process; thus, it is considered to be very fast. The above LAI methods were selected for benchmarking because FLARE, G-Nomix, and SALAI-NET are the newly published ones, reporting promising results; RFMix is the most popular and widely used for ancestry-related applications. Loter has the same problem formulation as Recomb-Mix does. HapMix was not included because it cannot tackle multi-way admixture and only produces inference results at the diploid level.

## Results

### Local ancestry inference for three-way inter-continental admixed populations

For three-way inter-continental simulated datasets, Recomb-Mix had the best *r*^2^ values and accuracy rates in reference panel sizes 100, 250, 500, and 1,000 with 15 generations and in generations 15, 50, 100, and 200 with 500 references. Figure 2 shows the *r*^2^ values of the inference results on six LAI methods, FLARE, G-Nomix, Loter, Recomb-Mix, RFMix, and SALAI-Net. Supplemental Figures S1 and S2 show the accuracy rates (values are in Supplemental Tables S5 and S6). Overall, as the reference panel size increases, the average *r*^2^ value and the accuracy rate increase for all methods. The large reference panel containing more individual samples than those in small ones helps improve the inference result. When using a reference panel with 1,000 individuals, all methods had at least 0.99 *r*^2^ value or 92% accuracy rate, while Recomb-Mix reached the best *r*^2^ value 0.9989 or accuracy rate of 99.10%.

**Figure 2:**
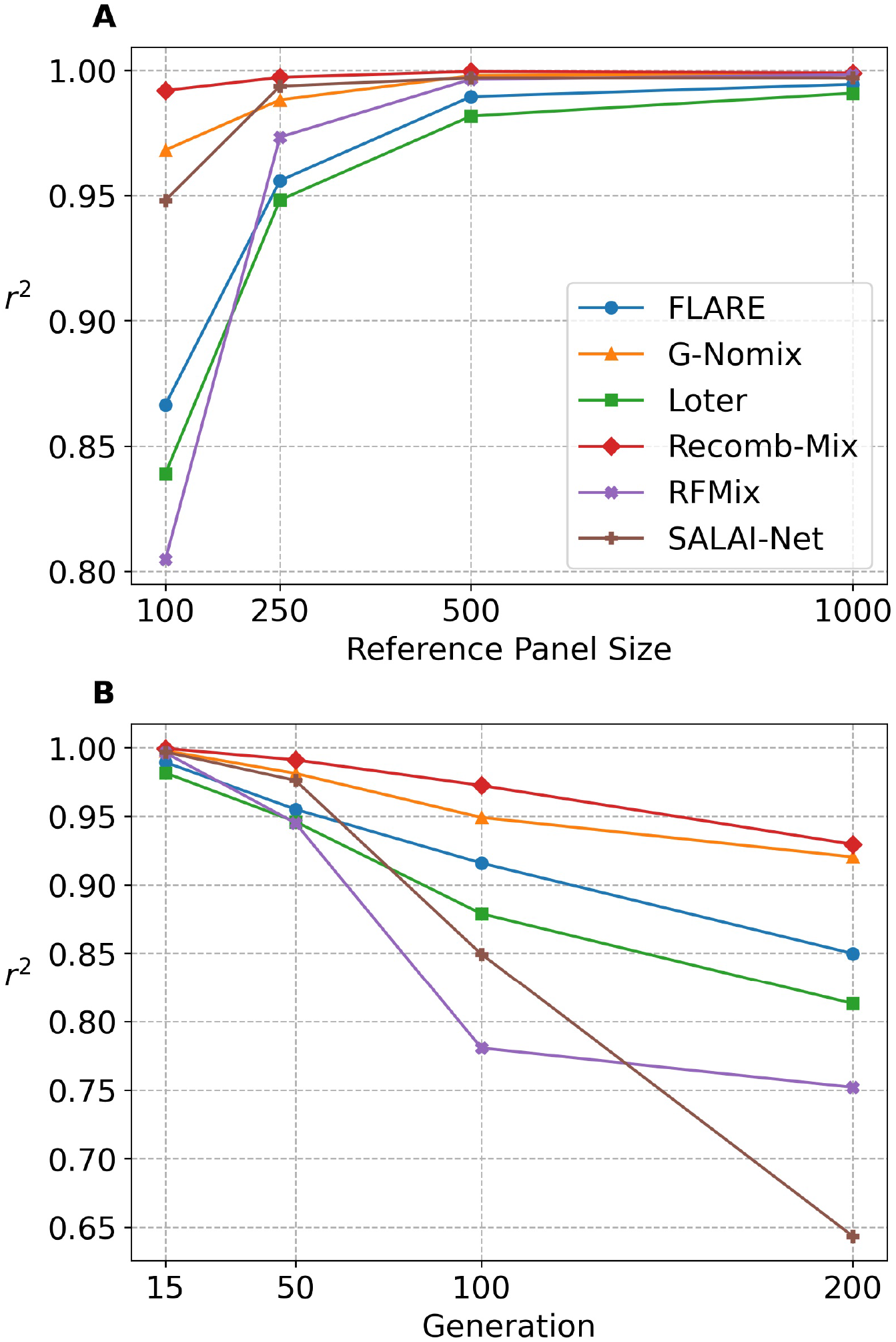
The squared Pearson’s correlation coefficient *r*^2^ of three-way inter-continental simulated datasets on FLARE, G-Nomix, Loter, Recomb-Mix, RFMix, and SALAI-Net. Markers were filtered with minor allele frequency ≤ 0.005 and minor allele count ≤ 50. (A) The three-way 15-generation datasets with the reference panel sizes 100, 250, 500, and 1,000 (values are in Supplemental Table S3). (B) The three-way 500-reference datasets with the generations 15, 50, 100, and 200 (values are in Supplemental Table S4).

Recomb-Mix performed well for a small reference panel (i.e., for a 100-individual penal it achieved the best performance, that is *r*^2^ value 0.9919 or accuracy rate of 97.96%). G-Nomix and SALAI-Net had the second and the third highest *r*^2^ values, which were 0.9681 and 0.9480. SALAI-Net and G-Nomix had the second and the third highest accuracy rates, which were 86.69% and 86.63%.

Recomb-Mix’s performance on ultra-small reference panels (i.e., size 20-50) was tested. Such cases are interesting because small reference panels can benefit low-resourced populations. Meanwhile, LAI with small reference panels is challenging because allele frequencies and haplotype frequencies are noisy. For a small reference panel size of 20, Recomb-Mix achieves the best accuracy rate of 62.85%; for a 50-individual reference panel, Recomb-Mix achieves 94.45%, while other methods’ accuracy rates are around 60% to 70% (see Supplemental Figure S3 and Table S7). We include MOSAIC (Salter-Townshend and Myers 2019) in this experiment as it reportedly performs well on small reference panels (Browning et al. 2023). MOSAIC achieves better accuracy rates than FLARE, Loter, and RFMix on reference panels of sizes 50 and 100, but its performance is worse than Recomb-Mix.

### Multi-way admixture

Besides the experiments on three-way admixed individuals, a case study on seven-way admixed individuals was investigated. The goal is to find out how LAI methods perform on individuals admixed from a large number of founder populations, as in a real case scenario, human individuals are involved in multiple population admixture events. Seven-way inter-continental datasets with various reference panel sizes and generations were simulated, and for such challenging datasets, the *r*^2^ values and the accuracy rates of LAI methods dropped, but Recomb-Mix kept performing well. Figure 3 shows the *r*^2^ values of the inference results on six LAI methods, FLARE, G-Nomix, Loter, Recomb-Mix, RFMix, and SALAI-Net. The average accuracy rates are in Supplemental Figures S4 and S5 (values are in Supplemental Tables S10 and S11). Figure 4B illustrates an inferred haplotype sample of an admixed individual from the methods. Compared to Figure 4A, more population labels were mistakenly assigned, as inferring seven-way admixed individuals is a harder task than inferring three-way ones. In general, window-based LAI methods performed better than site-based ones, except Recomb-Mix. Using a window as the smallest unit of inference helps tolerate errors within the window since the population label having the highest estimated probability determines the inference result for the entire window. On the other hand, site-based methods may focus more on a single site’s label, which may affect the inference result of one’s surrounding region when making incorrect inferences, especially if the number of potential population labels is large and the number of reference haplotype templates is limited. Though Recomb-Mix uses a site-based approach, it achieved high accuracy. Allowing individual variations within the same population helps inflate the panel so more reference haplotype templates (e.g., relatives) are available for local site inference.

**Figure 3:**
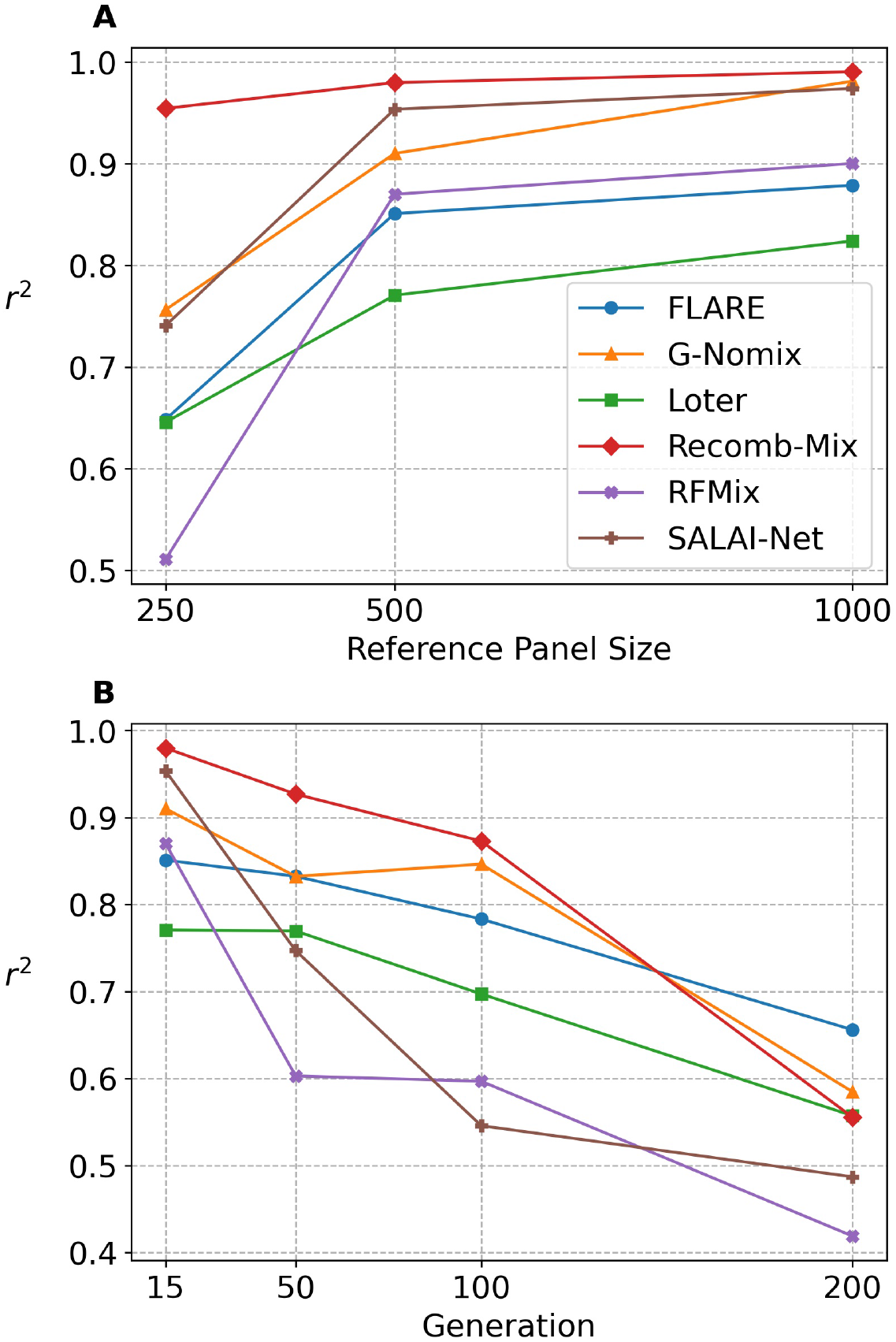
The squared Pearson’s correlation coefficient *r*^2^ of seven-way inter-continental simulated datasets on FLARE, G-Nomix, Loter, Recomb-Mix, RFMix, and SALAI-Net. Markers were filtered with minor allele frequency ≤ 0.005 and minor allele count ≤ 50. (A) The seven-way 15-generation datasets with the reference panel sizes 250, 500, and 1,000 (values are in Supplemental Table S8). The reference panel size 100 case was not included because the number of markers was too small and may have influenced the outcome after the filtering. (B) The seven-way 500-reference datasets with the generations 15, 50, 100, and 200 (values are in Supplemental Table S9).

**Figure 4:**
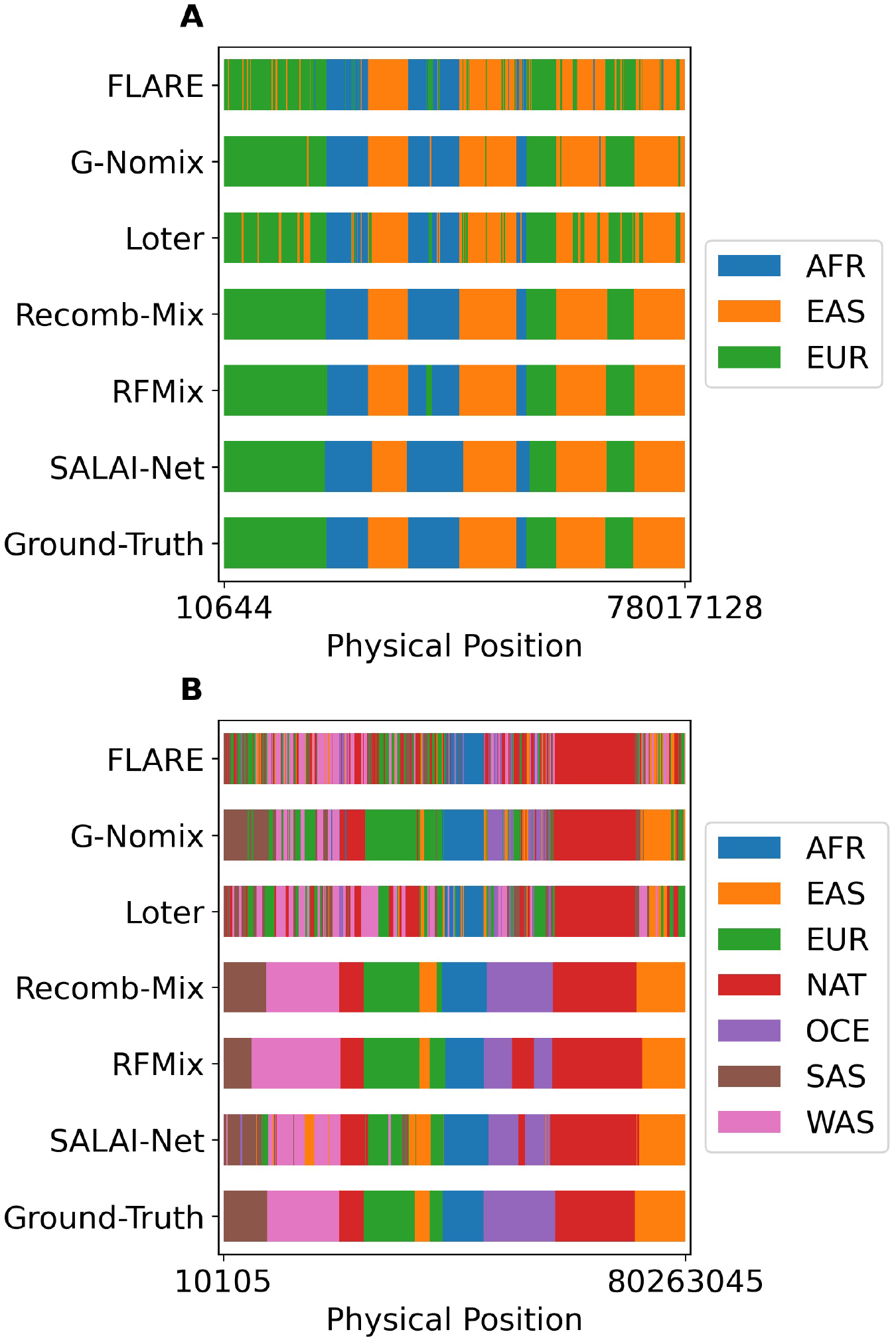
Sample haplotypes inferred by FLARE, G-Nomix, Loter, Recomb-Mix, RFMix, and SALAI-Net with the ground truth of ancestry labels. (A) An inferred sample haplotype from a three-way 15-generation 500-reference inter-continental simulated dataset. (B) An inferred sample haplotype from a seven-way 15-generation 500-reference inter-continental simulated dataset.

For a seven-way 200-generation 500-individual inter-continental WGS dataset, FLARE, G-Nomix, and Loter had better *r*^2^ values than Recomb-Mix. They were 0.1005, 0.0292, and 0.0017 higher than that of Recomb-Mix, respectively (see Figure 3B). FLARE, G-Nomix, and Loter are claimed to be the LAI methods good for identifying distant admixture events, as demonstrated high-resolution accuracy when the admixture event occurs over 100 generations (Browning et al. 2023; Hilmarsson et al. 2021; Dias-Alves et al. 2018). FLARE incorporates the number of generations as a parameter in their model and its value is updated using an iterative expectation maximization (EM) approach to calculate the probabilities of a change of ancestry state for each marker and haplotype. The longer the admixture event occurs, the higher the probability that ancestral segments or tracts having a large length difference appear. This information helps FLARE to update their generation parameter better. On top of G-Nomix’s base module’s classifier, a smoother module is added to refine the inference result. The smoother is a data-driven approach, which learns to capture the distribution of recombination breakpoints. Usually, the distant admixture event has richer information on the distribution of recombinations, which helps G-Nomix’s smoother module improve the accuracy. Loter adopts the bagging technique to generate the averaged result, which avoids putting a strong prior on a particular length of ancestry segment. This helps improve the inference accuracy since the ancestry segments appearing in distant admixture events are not the same length.

### Intra-continental admixture

Compared to the inter-continental admixture, the LAI on the intra-continental is relatively less studied. The same benchmarks were set up and evaluated as the inter-continental ones. Similar to the results of the inter-continental datasets, Recomb-Mix performed well on the intra-continental datasets. Figure 5A shows the *r*^2^ values of the local ancestry inference of six LAI methods, FLARE, G-Nomix, Loter, Recomb-Mix, RFMix, and SALAI-Net on a three-way 15-generation intra-continental simulated dataset. (Supplemental Figure S6 shows the average accuracy rates (values are in Supplemental Tables S14)). Overall, the *r*^2^ values of each method were worse than the ones in the inter-continental datasets. This is expected as the admixture occurring at the intra-continental level generates individuals who resemble each other. Thus, performing LAI on such datasets is more challenging than at the inter-continental level. Recomb-Mix had the best *r*^2^ value in reference panel sizes 250, 500, and 1,000 with 15 generations. For a 250-individual reference panel, the *r*^2^ value of Recomb-Mix was 0.9299, and the second-best method, G-Nomix, only achieved 0.8560. For a 1,000-individual reference panel, the *r*^2^ values of Recomb-Mix and G-Nomix were close (0.9800 and 0.9820).

**Figure 5:**
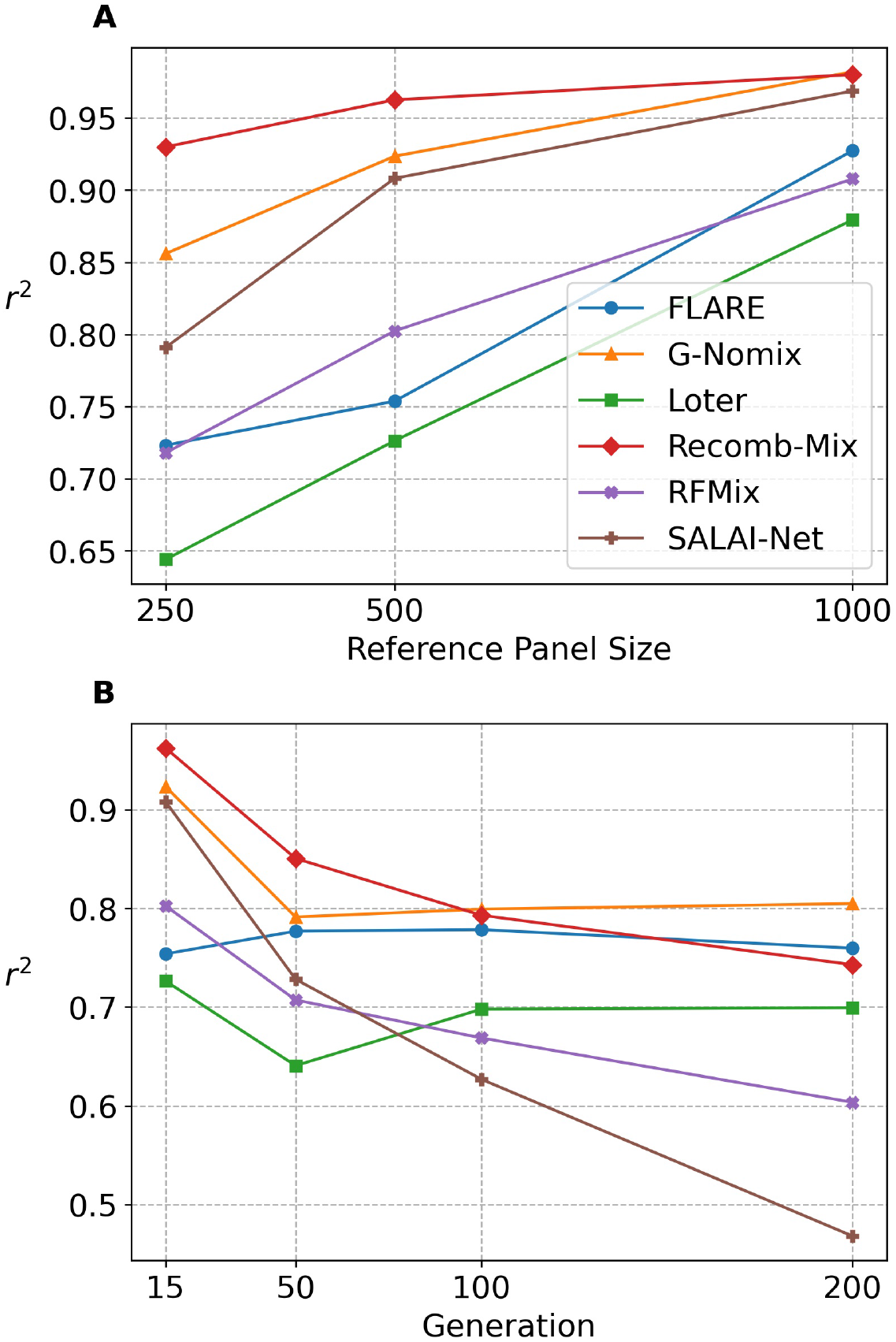
The squared Pearson’s correlation coefficient *r*^2^ of three-way intra-continental simulated datasets on FLARE, G-Nomix, Loter, Recomb-Mix, RFMix, and SALAI-Net. Markers were filtered with minor allele frequency ≤ 0.005 and minor allele count ≤ 50. (A) The three-way 15-generation datasets with the reference panel sizes 250, 500, and 1,000 (values are in Supplemental Table S12). The reference panel size 100 case was not included because the number of markers was too small and may have influenced the outcome after the filtering. (B) The three-way 500-reference datasets with the generations 15, 50, 100, and 200 (values are in Supplemental Table S13).

The impact of the number of generations on LAI at the intra-continental level was also investigated. Four three-way 500-reference intra-continental simulated datasets with generations 15, 50, 100, and 200 were tested, and the results show both the *r*^2^ values and accuracy rates are inversely proportional to the number of generations (see Figure 5B and Supplemental Figure S7 (values in Supplemental Table S15)). For a 15-generation dataset, Recome-Mix had the highest *r*^2^ value, 0.9625. The second and third high *r*^2^ values were 0.9235 (G-Nomix) and 0.9081 (SALAI-Net). For a 200-generation dataset, G-Nomix and FLARE had better *r*^2^ values than Recome-Mix. Also, Loter’s performance increased with the increasing number of generations. This result is consistent and observed in other simulated datasets, as G-Nomix, FLARE, and Loter do well in ancestry inference on the cases of distant admixture events.

### Robustness against admixture with uneven proportions of founders and references

To verify the robustness of Recomb-Mix handling cases on uneven founder populations and reference panels, LAI was experimented with using uneven founders for the imbalanced admixture simulation and uneven reference panels for the inference. Being uneven means the group consists of one-third of individuals from the first population, one-sixth from the second population, and half of individuals from the third population. Being even means the numbers of individuals from the populations in the group are divided equally. Three sets of experiments were performed. One three-way admixture dataset was simulated using even founders and inferred using even references, another was simulated using uneven founders and inferred using even references, and the other was simulated using even founders but inferred using uneven references.

The *r*^2^ values and the accuracy rates in Table 2 and Supplemental Table S16 indicate that admixed individuals with uneven founders and uneven reference panel slightly impact the performance across all LAI methods. Among all LAI methods, Recomb-Mix had the highest *r*^2^ values and accuracy rates in both cases (0.9426 or 89.20% and 0.8944 or 83.61%, respectively). The process of Recomb-Mix generating a collapsed graph helps convert the unbalanced reference populations into balanced ones. Thus, Recomb-Mix keeps a high accuracy of inference results on the unbalanced reference populations. Additionally, Recomb-Mix was tested on a modern Latino population admixture model that involves uneven founders, which is a popular realistic model used as a study case for the local ancestry inference (Maples et al. 2013; Wang et al. 2021). We used SLiM v4.0 to simulate the modern Latino population dataset on Chromosome 1, using the same settings from the RFMix paper (Maples et al. 2013). Ten Latino genomes with 45% Native American (NAT), 50% European (CEU), and 5% African ancestry (YRI) were simulated, originating from 400 individuals and 12 generations after the admixture event. 30 individuals from each population were used to form the reference panel. We used Beagle 5.4 to phase the source data. The average LAI accuracy rate using Recomb-Mix and RFMix was 99.36% and 93.79%, respectively. This shows that Recomb-Mix excels in the ancestry inference on the modern Latino population admixture model derived from the uneven founders.

**Table 2:** The squared Pearson’s correlation coefficient *r*^2^ of FLARE, G-Nomix, Loter, Recomb-Mix, RFMix, and SALAI-Net performing LAI on three-way 15-generation 500-reference intra-continental simulated datasets with even or uneven number of individuals per population in founder or reference panel. Markers were filtered with minor allele frequency ≤ 0.005 and minor allele count ≤ 50.

| Method | Even Founders<br>and References | Uneven Founders | Uneven<br>References |
| --- | --- | --- | --- |
| Loter | 0.7264 | 0.7263 | 0.7097 |
| FLARE | 0.7538 | 0.7187 | 0.7773 |
| RFMix | 0.8024 | 0.8112 | 0.7887 |
| SALAI-Net | 0.9081 | 0.8747 | 0.8692 |
| G-Nomix | 0.9235 | 0.9022 | 0.8811 |
| Recomb-Mix | **0.9625** | **0.9426** | **0.8944** |

### Robustness against ancestry misspecification panel

When performing real data analysis, the concern of data imperfection may be raised. Some populations may be less studied and underrepresented in available reference panels. Furthermore, the existing reference populations may contain a small fraction of admixture which may not make them the ideal proxies for the labeled populations. Thus, it is necessary to investigate the impact of the ancestry population misspecification on LAI.

An experiment was conducted by replacing the African reference population in a three-way inter-continental admixed dataset with an imperfect reference panel. The imperfect version of the African reference panel contains individuals who were Africans mixed with Europeans five generations from the start of the simulation. This approach has similar effects as the one MOSAIC had, where their imperfect reference panel contained admixed Sub-Saharan Africans and Europeans (Salter-Townshend and Myers 2019). We did not follow their process because the sampled individuals they used for the simulation were from the extended HGDP dataset, whose data density is only at the single nucleotide polymorphism (SNP) array level (Hellenthal et al. 2014). The ancestry misspecification experiments were repeated for 15, 50, 100, and 200 generations since the admixture event, and FLARE, G-Nomix, Loter, Recomb-Mix, RFMix, and SALAI-Net were tested. We did not include MOSAIC as it was designed for the case when the source population lacked the availability of WGS data (Salter-Townshend and Myers 2019).

All LAI methods were impacted by the misspecification reference panel but still performed well, as shown in Figure 6. Under the *r*^2^ criteria with markers having minor allele values filtered, Recomb-Mix performed the best in the cases of 50 and 100 generations since the admixture event. RFMix performed the best for the most recent admixture case, and FLARE performed the best for the most distant admixture case. Without filtering out any markers, Recomb-Mix had the highest accuracy rate for most cases except the 15-generation case where RFMix performed the best.

**Figure 6:**
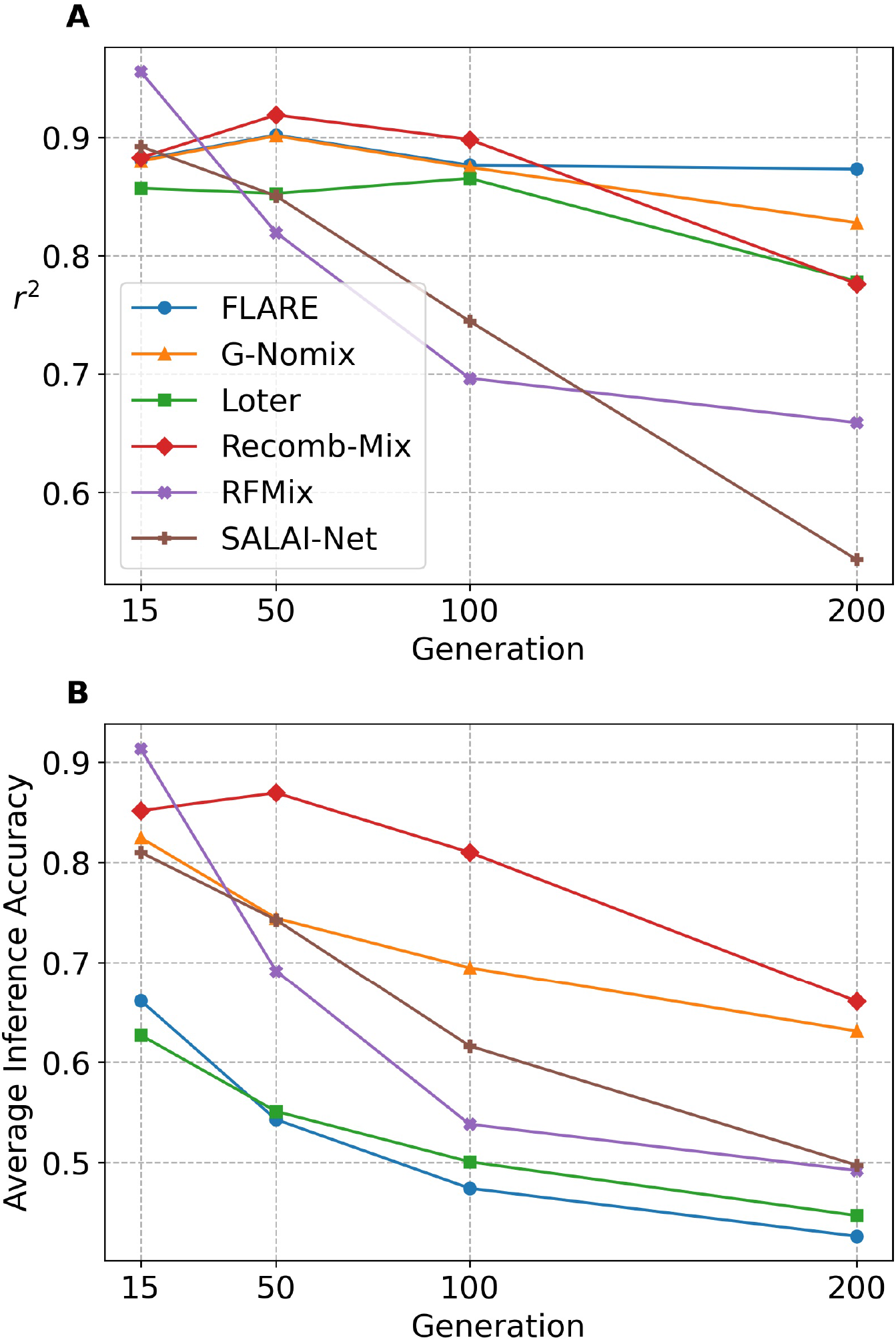
The performance of local ancestry inference with generations 15, 50, 100, and 200 of the three-way 500-misspecified-reference inter-continental simulated datasets on FLARE, G-Nomix, Loter, Recomb-Mix, RFMix, and SALAI-Net. (A) The squared Pearson’s correlation coefficient *r*^2^ (values are in Supplemental Table S17). Markers were filtered with minor allele frequency ≤ 0.005 and minor allele count ≤ 50. (B) The average accuracy rates (values are in Supplemental Table S18).

### Robustness against phasing error

To investigate the impact of phasing error on local ancestry inference, two cases of phasing errors on either the target panel or the reference panel were explored. The three-way 15-generation 100-reference inter-continental simulated dataset was used as the baseline, and Beagle 5.4 was applied to phase the panels. The phasing error rate was 0.58% for the target panel and 1.33% for the reference panel, verified by VCFtools (Danecek et al. 2011). FLARE, G-Nomix, Loter, Recomb-Mix, RFMix, and SALAI-Net were tested on the panels and diploid accuracy rates were used for the performance measurement as RFMix did (Maples et al. 2013). The results in Supplemental Table S19 show that the diploid accuracy rates did not fluctuate much when there were phasing errors on the panels, indicating that the low rate of phasing errors may not have a substantial impact on the local ancestry inference.

### Recomb-Mix is efficient in memory, space, and run time

We examined the run time and maximum amount of memory LAI methods used for their performance on admixed individual haplotypes. Supplemental Figures S8, S9, and Table S20 show the average CPU run time and maximum amount of physical memory that all six LAI methods consumed across different experimental runs. In general, for the same method, inference on three-way admixed individuals was faster than those on seven-way. This is expected as a seven-way admixture has many more local ancestral segments across the chromosome than ones in a three-way, which costs more time for the inference. All methods showed reasonable run time for an LAI query of an admixed individual haplotype except Loter, which was about 10 or 100 times slower than other methods. SALAI-Net was the fastest method and Recomb-Mix was the runner-up but only took 0.31 and 2.04 more seconds than SALAI-Net in three-way and seven-way datasets. From the memory-consuming perspective, all methods’ memory usage was acceptable, and Recomb-Mix required the smallest amount of memory, 2.44 and 4.13 GB in three-way and seven-way datasets, respectively.

Recomb-Mix has a feature that converts the compact population graph into a Variant Call Format (VCF) file (Danecek et al. 2011). Later, they can be reused by Recomb-Mix to save processing time. A compact VCF file is much smaller than the original one since it only contains individual haplotype templates with population-level information. For example, the disk space needed to store a 3-way inter-continental 500-reference panel was decreased from 665 to 13.3 MB (and 1.4 MB for a compressed VCF file). Similarly, for a 7-way panel, the disk space was decreased from 787 to 21.8 MB (and 1.9 MB for a compressed VCF file).

### The 1000 Genomes Project and the Human Genome Diversity Project ancestry analysis

To show the scalability and robustness of Recomb-Mix, we estimated the ancestry proportions from the inferred local ancestries for the populations in the 1000 Genomes Project (TGP) data (Byrska-Bishop et al. 2022) using the four founder populations (Africans, Admixed Americans, East Asians, and Europeans) as the reference panel from the Human Genome Diversity Project (HGDP) data (Bergström et al. 2020). Similarly, We estimated the ancestry proportions for the populations in the HGDP data using the four founder populations (Africans, East Asians, Europeans, and Native Americans) from the TGP data. We merged two Chromosome 18 datasets (TGP with 3,457,645 markers and HGDP with 2,127,412 markers), yielding 1,165,399 intersected markers. Then the merged dataset was phased using Beagle 5.4 and the individuals were assigned the population labels provided by their original datasets. Like FLARE (Browning et al. 2023), we calculated the global ancestry composition by averaging the estimated local ancestry proportions across the genome. Figure 7 is Recomb-Mix’s ancestry inference result on the TGP dataset that is generally consistent with exceptions. Similar results using other LAI methods are available in Supplemental Figure S10. For African individuals who reside in the African continent, at least 97% segment was labeled as African on average. For ACB and ASW individuals (located in the Caribbean and America), a small portion of the segment was labeled as non-African due to their admixed backgrounds. Most segments of American individuals were labeled as mixed percentages of Europeans and Native Americans. South Asian individuals were labeled as mixed percentages of East Asian and European. East Asian and European individuals had at least 99% and 94% segments labeled as East Asian and European, respectively.

**Figure 7:**
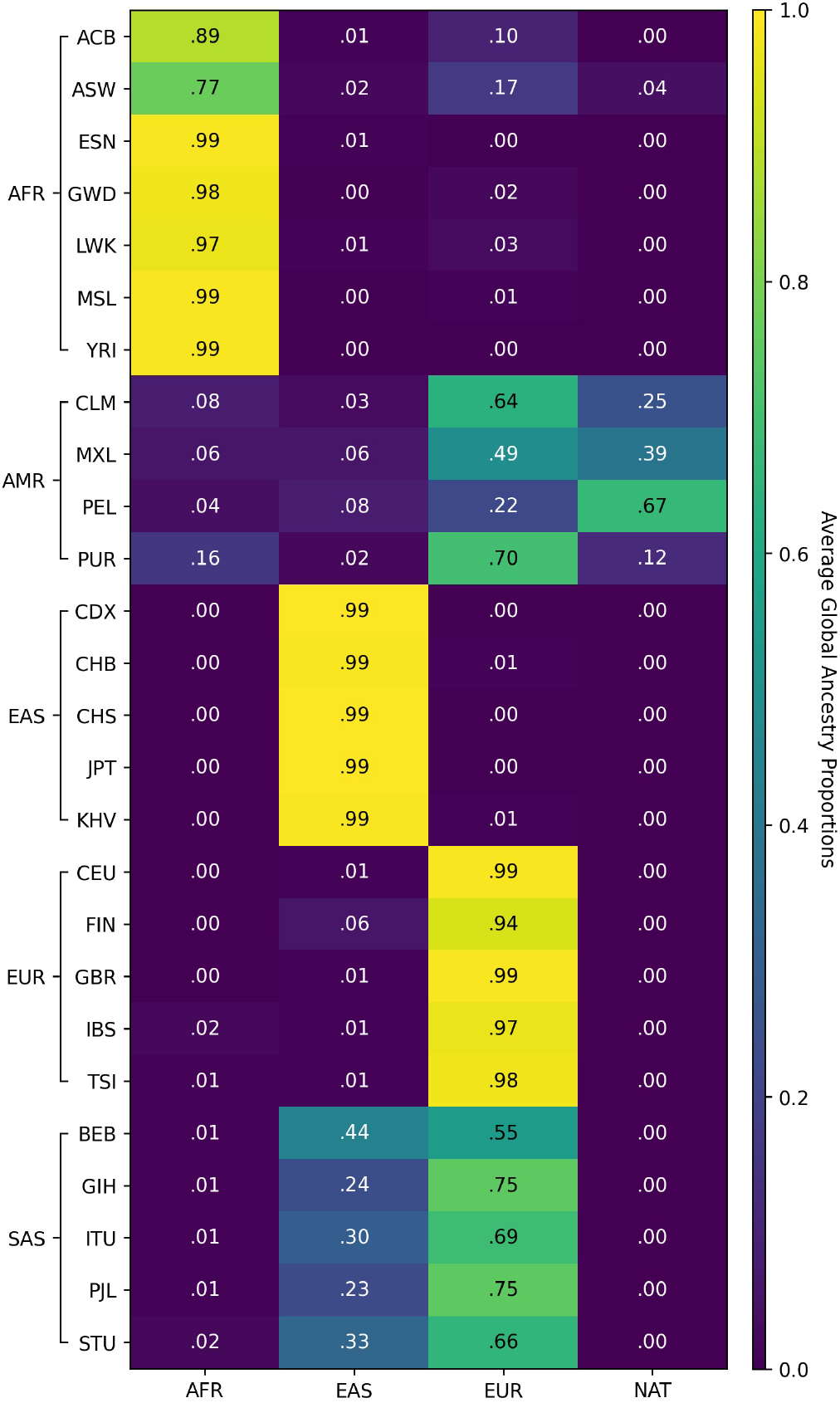
The average global ancestry proportions in the TGP Chromosome 18 data using four reference ancestries from the HGDP data. Descriptions of the populations are in Supplemental Table S1.

The ancestry inference result on the HGDP dataset is generally anticipated. Figure 8 shows that Recomb-Mix predicted African, East Asian, European, and native American individuals have 99%, 94%, 88%, and 98% segments matched expected ancestries. For the Oceanian individuals, the segments were decomposed into mixed ancestries, primarily East Asian and African. For South Asian and West Asian individuals, the segments were inferred as mixed and mainly consisted of European ancestry. Interpretation of ancestry inference results when the founder populations in the reference panel were more complex. For example, recent genetic evidence suggests that EUR, WAS, and SAS may share some Yamnaya DNA (Lazaridis et al. 2022; Narasimhan et al. 2019), which might be part of the causes of our results of possible shared (about 10%) ancient ancestry among AMR, EUR, OCE, SAS, and WAS. Similar behaviors were observed on other LAI methods, such as G-Nomix and SALAI-Net (see Supplemental Figure S11). Recomb-Mix was forced to give a single LAI call for these regions since the inference results were based on the given reference panels. If the ancient population were not in the reference panel, the segment would be labeled as the population closest to the ancient one.

**Figure 8:**
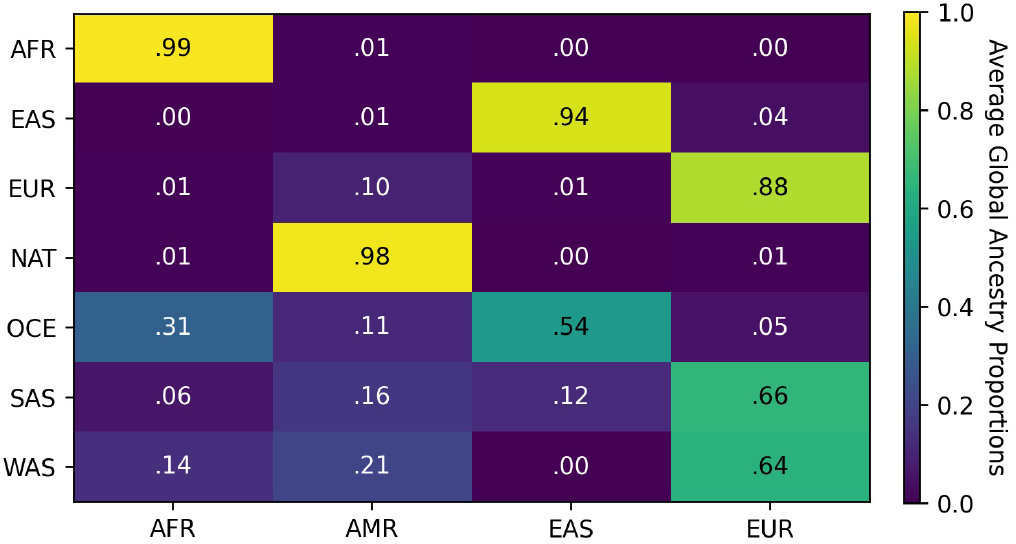
The average global ancestry proportions in the HGDP Chromosome 18 data using four reference ancestries from the TGP data. Descriptions of the populations are in Supplemental Table S1.

### Discrete ancestry informative markers

Local ancestry inference may benefit from ancestry informative markers (AIMs), which are genetic markers with significantly different allele frequencies in various populations (Parra et al. 1998). AIMs provide information regarding ancestry and can be determined in a panel by measuring marker informativeness for ancestry (Ding et al. 2011). Rather than relying on a selected set of markers, i.e., AIMs, the proposed compact population graph keeps all markers, and the only differential information between populations is that some alleles might be missing in one or several populations. These markers are dubbed discrete AIMs (dAIMs), whose allele values in one population that at least one of the other populations does not have. dAIMs are operationally defined and depend on some random chance of whether an allele is present in the reference panel for the population or not. However, as it is shown in Figure 9 (the dAIM densities (i.e., percentages of markers in the dataset being dAIMs) of Chromosome 18 in the HGDP data (Bergström et al. 2020)), dAIMs are densely available on a typical panel. Therefore, even though collapsing nodes reduced the information in the original panel, the remaining information in dAIMs might be sufficient for making good-quality ancestry calls. For example, there is a dAIM density peak occurring around the 18q21 region in the HGDP dataset (see Figure 9). In a previous admixture mapping study, a genome-wide significant admixture mapping peak contributed from multiple ancestry signals was identified in the same region (Gignoux et al. 2019). This correlation suggests that dAIM density has the potential to identify ancestry-specific selection.

**Figure 9:**
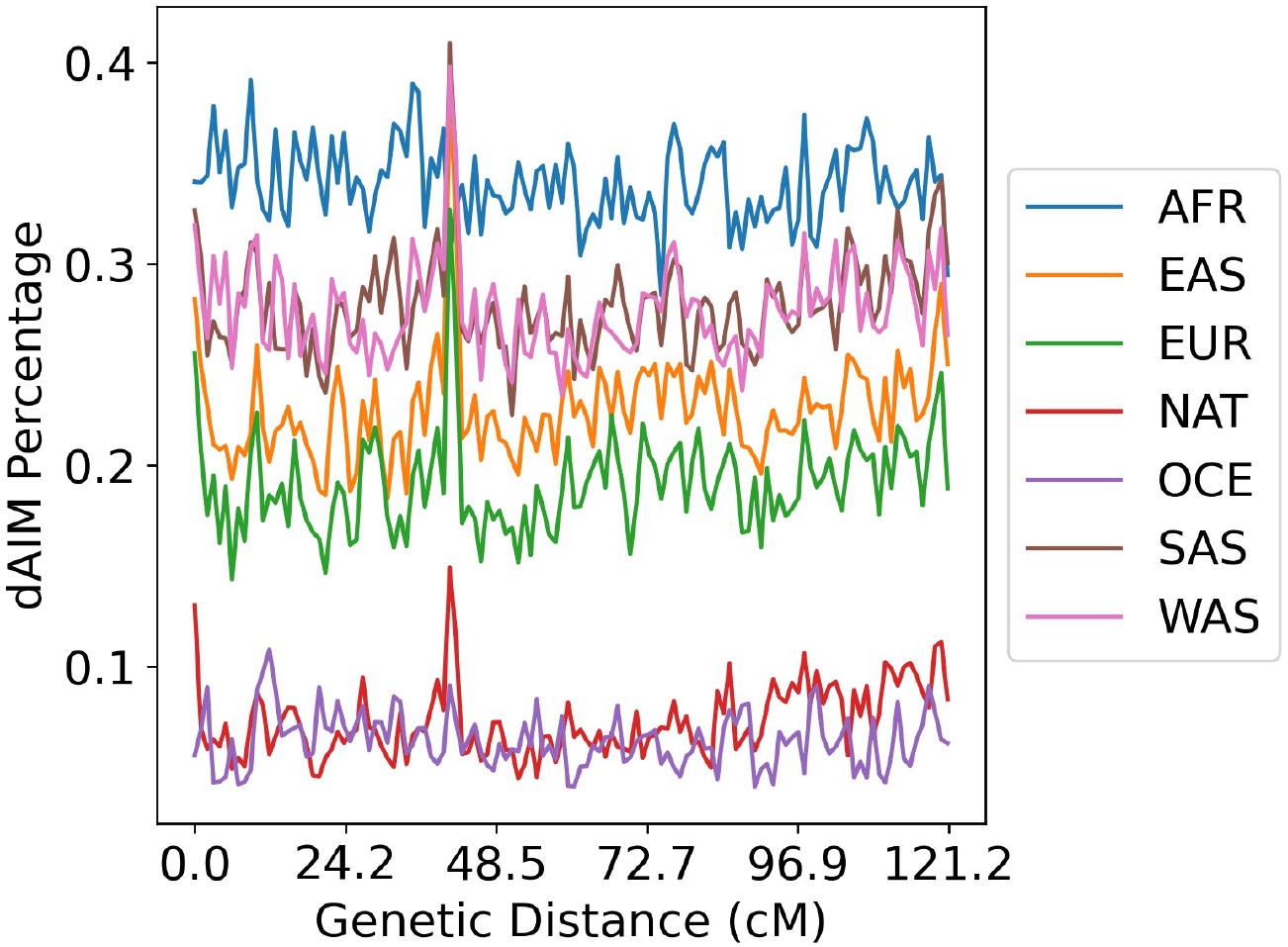
The discrete AIM (dAIM) density in the HGDP dataset per population on Chromosome 18. Each bin is 1 centiMorgan (cM), showing the markers’ dAIM percentage.

### Ablation study

We want to understand which component contributes the most to Recomb-Mix’s ancestry inference process. Experiments were designed for Recomb-Mix to make inferences on a three-way inter-continental 15-generation 100-reference simulated dataset by not setting the within-population template change penalty to zero or not using the recombination rates. We observed a slight decrease in performance when the recombination rate was not used. However, the performance dropped significantly when the within-population template change penalty was applied (see Supplemental Figure S12 and Table S21). Supplemental Figure S12 shows LAI accuracies are significantly improved when setting the within-population template change penalty to zero, especially for small reference panel cases. We also calculated the average number of threading path changes across populations and the standard deviations of the dataset with 228, 503 markers for Recomb-Mix and the version that used the within-population template change penalty. Recomb-Mix had 8.47 ± 2.54, while the latter version had a much larger number, 198.46 ± 46.94. The LAI calling became less effective when using the template change penalty within each population. This may be due to the numerous local optimal threading paths to be explored within each population, which can lead to noise and deviation from finding the path with the minimal global penalty score. By setting the within-population template change penalty to zero, the number of explorations between the paths within a population is significantly reduced, and the focus is shifted to only a few consolidated paths representing diverse haplotype templates.

The dAIMs are usually evenly distributed alongside the chromosome, as we showed in Figure 9.

To illustrate dAIM’s important role in LAI, we designed an experiment to engineer a new dataset based on the simulated one by taking out all the dAIMs for certain regions. We tested Recomb-Mix on the engineered dataset, and the result showed a strong correlation between the dAIM density and the accuracy of the inference. Supplemental Figures S13 and S14 show the dAIM density and local ancestry inference accuracy rate of the original dataset and the engineered dataset. In Supplemental Figure S14, there are five instances where the low LAI accurate rates correspond with areas lacking dAIMs in the engineered dataset. The Pearson correlation coefficient for this dataset’s dAIM density and local ancestry inference accuracy rate is 0.79, demonstrating a strong correlation between dAIMs and LAI accuracies.

### Single nucleotide polymorphism array data analysis

We created an SNP array dataset to verify if Recomb-Mix works on a panel with a limited number of dAIMs and rare variants. First, we followed Tang et al.’s pipeline (2022) to down-sample the three-way inter-continental 15-generation 100-reference simulated sequencing dataset. The number of markers was decreased from 228, 503 to 15, 584. Then, we further filtered out markers having minor allele frequency (MAF) less than 5%, as typically genotyping array data contains common variants whose MAF *>* 5% (Bomba et al. 2017; Verlouw et al. 2021). Eventually, the dataset had 8, 744 markers that resembled an imputed SNP array panel. The number of dAIMs in the panel also decreased compared to the one in the sequencing panel (see Supplemental Figure S15 and S16); however, the dAIM density in the SNP array data did not change as much as the one in the sequencing data (see Supplemental Figure S17 and S18). We tested Recomb-Mix on the SNP array dataset, and it achieved 0.9949 *r*^2^ value and 96.34% accuracy rate, compared to 0.9986 and 97.96% on the sequencing dataset without filtering any sites. This result suggests Recomb-Mix has the same performance on the SNP array data as on the sequencing data, possibly due to the dAIM density in the SNP array data being on the same order of magnitude as the one in the sequencing data.

## Discussion

We presented a new LAI method named Recomb-Mix, based on a simplified LS model formulating LAI as a graph optimization problem. By not considering recombination penalties within populations, Recomb-Mix shows promising LAI results under various circumstances. A compact population graph also helps Recomb-Mix process LAI effectively and efficiently. Furthermore, it is convenient to store the reference panel as a compact population graph on disk, which takes up little space for future ancestry inference without a re-transformation process. Recomb-Mix is competitive with other state-of-the-art LAI methods in accuracy and computational performance and is applicable to real genomic datasets. We introduced the concept of dAIM, where dAIMs are determined by the allele values of each population. We showed that dAIMs can have marker informativeness for ancestry. Of course, this selection of markers is simplistic and mainly captures the differentially present or absent markers in the reference panel across populations. In future studies, other ways of collapsing the graph might be explored to retain more relevant ancestry information from the reference panel. It might be optimized to allow the selection of markers present in multiple populations but with different allele frequencies. This could further enhance the performance and enable our model to provide uncertainty estimates for ancestry inference results.

As a site-based LAI method, Recomb-Mix is designed to exploit the site-level information to achieve superior accuracy, especially at the intra-continental level. The number of dAIMs in intra-continental admixed individuals is less than those in inter-continental admixed individuals, as intra-dAIMs are a subset of inter-dAIMs. Window-based LAI methods may find it difficult to achieve high accuracy during the intra-level ancestry inference. There is a higher probability for each window containing multiple intra-dAIMs than that for inter-dAIMs. Since the window is the smallest unit representing one ancestral source, windows having intra-dAIMs representing different populations may easily misrepresent the inference result. Decreasing the window size may mitigate the situation, with the potential computational burden. However, its lower bound is one site per window, i.e., site-based.

Despite the high accuracy rates demonstrated, Recomb-Mix has limitations of not considering the disparate genetic maps across populations, the allele frequencies, or genotyping and phasing errors. Thus, other complementary methods may be useful for a well-specified model. FLARE takes optional parameters such as minor allele frequency and number of generations since admixture. It may perform well if these biological parameters are correctly estimated for the model. G-Nomix has a few pre-trained models available which may be in handy if one was pre-trained specifically for the given model. SALAI-Net may be a good choice as it employs a pre-trained model that is generalized and applicable to any species and any set of ancestries. Though Recomb-Mix was not designed to handle erroneous panels, the genotyping error and phasing error seem to have no large impact on the inference results (see the Results section). If the error rate is high, a pre-processing step may be needed to correct the noisy data panel before making the inference.

## Software availability

The Recomb-Mix code is available at https://github.com/ucfcbb/Recomb-Mix.

## Competing interest statement

The authors declare no competing interests.

## Acknowledgments

The authors thank Dr. Ahsan Sanaullah for the helpful suggestions on the manuscript. This work was supported by the National Institutes of Health under grants R01HG010086 and R56HG011509.

## Notes

### Competing Interest Statement

The authors have declared no competing interest.

### Summary of Updates

Overall text was revised. Problem formulation was amended. New results were included.

## References

1. Adrion JR, Cole CB, Dukler N, Galloway JG, Gladstein AL, Gower G, Kyriazis CC, Ragsdale AP, Tsambos G, Baumdicker F, et al.. 2020. A community-maintained standard library of population genetic models. eLife 9: e54967.

2. Atkinson EG, Maihofer AX, Kanai M, Martin AR, Karczewski KJ, Santoro ML, Ulirsch JC, Kamatani Y, Okada Y, Finucane HK, et al.. 2021. Tractor uses local ancestry to enable the inclusion of admixed individuals in GWAS and to boost power. Nature Genetics 53: 195–204.

3. Auton A, Abecasis GR, Altshuler DM, Durbin RM, Bentley DR, Chakravarti A, Clark AG, Donnelly P, Eichler EE, Flicek P, et al.. 2015. A global reference for human genetic variation. Nature 526: 68–74.

4. Baran Y, Pasaniuc B, Sankararaman S, Torgerson DG, Gignoux C, Eng C, Rodriguez-Cintron W, Chapela R, Ford JG, Avila PC, et al.. 2012. Fast and accurate inference of local ancestry in Latino populations. Bioinformatics 28: 1359–1367.

5. Bergström A, McCarthy SA, Hui R, Almarri MA, Ayub Q, Danecek P, Chen Y, Felkel S, Hallast P, Kamm J, et al.. 2020. Insights into human genetic variation and population history from 929 diverse genomes. Science 367: eaay5012.

6. Bomba L, Walter K, and Soranzo N. 2017. The impact of rare and low-frequency genetic variants in common disease. Genome Biology 18: 77.

7. Browning BL, Tian X, Zhou Y, and Browning SR. 2021. Fast two-stage phasing of large-scale sequence data. The American Journal of Human Genetics 108: 1880–1890.

8. Browning BL, Zhou Y, and Browning SR. 2018. A One-Penny imputed genome from Next-Generation reference panels. The American Journal of Human Genetics 103: 338–348.

9. Browning SR, Waples RK, and Browning BL. 2023. Fast, accurate local ancestry inference with FLARE. The American Journal of Human Genetics 110: 326–335.

10. Bycroft C, Freeman C, Petkova D, Band G, Elliott LT, Sharp K, Motyer A, Vukcevic D, Delaneau O, O’Connell J, et al.. 2018. The UK Biobank resource with deep phenotyping and genomic data. Nature 562: 203–209.

11. Byrska-Bishop M, Evani US, Zhao X, Basile AO, Abel HJ, Regier AA, Corvelo A, Clarke WE, Musunuri R, Nagulapalli K, et al.. 2022. High-coverage whole-genome sequencing of the expanded 1000 Genomes Project cohort including 602 trios. Cell 185: 3426–3440.e19.

12. Church DM, Schneider VA, Graves T, Auger K, Cunningham F, Bouk N, Chen HC, Agarwala R, McLaren WM, Ritchie GR, et al.. 2011. Modernizing reference genome assemblies. PLoS Biology 9: 1–5.

13. Clarke L, Fairley S, Zheng-Bradley X, Streeter I, Perry E, Lowy E, Tasśe AM, and Flicek P. 2016. The international Genome sample resource (IGSR): A worldwide collection of genome variation incorporating the 1000 Genomes Project data. Nucleic Acids Research 45: D854–D859.

14. Danecek P, Auton A, Abecasis G, Albers CA, Banks E, DePristo MA, Handsaker RE, Lunter G, Marth GT, Sherry ST, et al.. 2011. The variant call format and VCFtools. Bioinformatics 27: 2156–2158.

15. Delaneau O, Marchini J, and Zagury JF. 2012. A linear complexity phasing method for thousands of genomes. Nature Methods 9: 179–181.

16. Delaneau O, Zagury JF, Robinson MR, Marchini JL, and Dermitzakis ET. 2019. Accurate, scalable and integrative haplotype estimation. Nature Communications 10: 5436.

17. Dias-Alves T, Mairal J, and Blum MGB. 2018. Loter: A software package to infer local ancestry for a wide range of species. Molecular Biology and Evolution 35: 2318–2326.

18. Ding L, Wiener H, Abebe T, Altaye M, Go RC, Kercsmar C, Grabowski G, Martin LJ, Khurana Hershey GK, Chakorborty R, et al.. 2011. Comparison of measures of marker informativeness for ancestry and admixture mapping. BMC Genomics 12: 622.

19. Ding Y, Hou K, Xu Z, Pimplaskar A, Petter E, Boulier K, Privé F, Vilhjälmsson BJ, Olde Loohuis LM, and Pasaniuc B. 2023. Polygenic scoring accuracy varies across the genetic ancestry continuum. Nature 618: 774–781.

20. Duncan L, Shen H, Gelaye B, Meijsen J, Ressler K, Feldman M, Peterson R, and Domingue B. 2019. Analysis of polygenic risk score usage and performance in diverse human populations. Nature Communications 10: 3328.

21. Durand EY, Do CB, Wilton PR, Mountain JL, Auton A, Poznik GD, and Macpherson JM. 2021. A scalable pipeline for local ancestry inference using tens of thousands of reference haplotypes. bioRxiv doi: 10.1101/2021.01.19.427308.

22. Durbin R. 2014. Efficient haplotype matching and storage using the positional Burrows-Wheeler transform (PBWT). Bioinformatics 30: 1266–1272.

23. Geza E, Mugo J, Mulder NJ, Wonkam A, Chimusa ER, and Mazandu GK. 2018. A comprehensive survey of models for dissecting local ancestry deconvolution in human genome. Briefings in Bioinformatics 20: 1709–1724.

24. Gignoux CR, Torgerson DG, Pino-Yanes M, Uricchio LH, Galanter J, Roth LA, Eng C, Hu D, Nguyen EA, Huntsman S, et al.. 2019. An admixture mapping meta-analysis implicates genetic variation at 18q21 with asthma susceptibility in Latinos. Journal of Allergy and Clinical Immunology 143: 957–969.

25. Haller BC, Galloway J, Kelleher J, Messer PW, and Ralph PL. 2019. Tree-sequence recording in SLiM opens new horizons for forward-time simulation of whole genomes. Molecular Ecology Resources 19: 552–566.

26. Haller BC and Messer PW. 2019. SLiM 3: Forward genetic simulations beyond the Wright–Fisher model. Molecular Biology and Evolution 36: 632–637.

27. Hamid I, Korunes KL, Schrider DR, and Goldberg A. 2023. Localizing post-admixture adaptive variants with object detection on ancestry-painted chromosomes. Molecular Biology and Evolution 40: msad074.

28. Hellenthal G, Busby GBJ, Band G, Wilson JF, Capelli C, Falush D, and Myers S. 2014. A genetic atlas of human admixture history. Science 343: 747–751.

29. Hilmarsson H, Kumar AS, Rastogi R, Bustamante CD, Mas Montserrat D, and Ioannidis AG. 2021.High resolution ancestry deconvolution for next generation genomic data. bioRxiv doi: 10.1101/2021.09.19.460980.

30. Hou K, Ding Y, Xu Z, Wu Y, Bhattacharya A, Mester R, Belbin GM, Buyske S, Conti DV, Darst BF, et al.. 2023. Causal effects on complex traits are similar for common variants across segments of different continental ancestries within admixed individuals. Nature Genetics 55: 549–558.

31. Kurki MI, Karjalainen J, Palta P, Sipilä TP, Kristiansson K, Donner KM, Reeve MP, Laivuori H, Aavikko M, Kaunisto MA, et al. 2023. Finngen provides genetic insights from a well-phenotyped isolated population. Nature 613: 508–518.

32. Lawson DJ, Hellenthal G, Myers S, and Falush D. 2012. Inference of population structure using dense haplotype data. PLoS Genetics 8: 1–16.

33. Lazaridis I, Alpaslan-Roodenberg S, Acar A, Açıkkol A, Agelarakis A, Aghikyan L, Akyüz U, Andreeva D, Andrijašević G, Antonović D, et al.. 2022. The genetic history of the Southern Arc: A bridge between West Asia and Europe. Science 377: eabm4247.

34. Li N and Stephens M. 2003. Modeling linkage disequilibrium and identifying recombination hotspots using single-nucleotide polymorphism data. Genetics 165: 2213–2233.

35. Maples BK, Gravel S, Kenny EE, and Bustamante CD. 2013. RFMix: A discriminative modeling approach for rapid and robust local-ancestry inference. The American Journal of Human Genetics 93: 278–288.

36. Martin AR, Gignoux CR, Walters RK, Wojcik GL, Neale BM, Gravel S, Daly MJ, Bustamante CD, and Kenny EE. 2017. Human demographic history impacts genetic risk prediction across diverse populations. The American Journal of Human Genetics 100: 635–649.

37. Montserrat DM, Bustamante C, and Ioannidis A. 2020. Lai-Net: Local-ancestry inference with neural networks. In ICASSP 2020 - 2020 IEEE International Conference on Acoustics, Speech and Signal Processing (ICASSP), pp. 1314–1318.

38. Narasimhan VM, Patterson N, Moorjani P, Rohland N, Bernardos R, Mallick S, Lazaridis I, Nakatsuka N, Olalde I, Lipson M, et al.. 2019. The formation of human populations in South and Central Asia. Science 365: eaat7487.

39. Oriol Sabat B, Mas Montserrat D, Giro-i-Nieto X, and Ioannidis AG. 2022. SALAI-Net: Species-agnostic local ancestry inference network. Bioinformatics 38: ii27–ii33.

40. Parra EJ, Marcini A, Akey J, Martinson J, Batzer MA, Cooper R, Forrester T, Allison DB, Deka R, Ferrell RE, et al.. 1998. Estimating African American admixture proportions by use of population-specific alleles. The American Journal of Human Genetics 63: 1839–1851.

41. Pasaniuc B, Zaitlen N, Lettre G, Chen GK, Tandon A, Kao WHL, Ruczinski I, Fornage M, Siscovick DS, Zhu X, et al.. 2011. Enhanced statistical tests for GWAS in admixed populations: Assessment using African Americans from CARe and a breast cancer consortium. PLoS Genetics 7: 1–15.

42. Patin E, Siddle KJ, Laval G, Quach H, Harmant C, Becker N, Froment A, Ŕegnault B, Lemée L, Gravel S, et al.. 2014. The impact of agricultural emergence on the genetic history of african rainforest hunter-gatherers and agriculturalists. Nature Communications 5: 3163.

43. Price AL, Tandon A, Patterson N, Barnes KC, Rafaels N, Ruczinski I, Beaty TH, Mathias R, Reich D, and Myers S. 2009. Sensitive detection of chromosomal segments of distinct ancestry in admixed populations. PLoS Genetics 5: 1–18.

44. Reich D, Patterson N, Jager PLD, McDonald GJ, Waliszewska A, Tandon A, Lincoln RR, DeLoa C, Fruhan SA, Cabre P, et al.. 2005. A whole-genome admixture scan finds a candidate locus for multiple sclerosis susceptibility. Nature Genetics 37: 1113–1118.

45. Salter-Townshend M and Myers S. 2019. Fine-scale inference of ancestry segments without prior knowledge of admixing groups. Genetics 212: 869–889.

46. Schneider VA, Graves-Lindsay T, Howe K, Bouk N, Chen HC, Kitts PA, Murphy TD, Pruitt KD, Thibaud-Nissen F, Albracht D, et al.. 2017. Evaluation of GRCh38 and de novo haploid genome assemblies demonstrates the enduring quality of the reference assembly. Genome Research 27: 849– 864.

47. Suarez-Pajes E, Díaz-de Usera A, Marcelino-Rodŕıguez I, Guillen-Guio B, and Flores C. 2021. Genetic Ancestry Inference and Its Application for the Genetic Mapping of Human Diseases. International Journal of Molecular Sciences 22: 6962.

48. Tang K, Naseri A, Wei Y, Zhang S, and Zhi D. 2022. Open-source benchmarking of IBD segment detection methods for biobank-scale cohorts. GigaScience 11. Giac111.

49. Verlouw JAM, Clemens E, de Vries JH, Zolk O, Verkerk AJMH, am Zehnhoff-Dinnesen A, Medina-Gomez C, Lanvers-Kaminsky C, Rivadeneira F, Langer T, et al.. 2021. A comparison of genotyping arrays. European Journal of Human Genetics 29: 1611–1624.

50. Wang Y, Song S, Schraiber JG, Sedghifar A, Byrnes JK, Turissini DA, Hong EL, Ball CA, and Noto K. 2021. Ancestry inference using reference labeled clusters of haplotypes. BMC Bioinformatics 22: 459.

51. Wu J, Liu Y, and Zhao Y. 2021. Systematic review on local ancestor inference from a mathematical and algorithmic perspective. Frontiers in Genetics 12: 698.

