## Supplemental Material for "Fast and accurate local ancestry inference with Recomb-Mix"

| Population Code | Population Description |
|---|---|
| ACB | African Caribbeans in Barbados |
| AFR | African |
| AMR | Admixed American |
| ASW | Americans of African Ancestry in SW USA |
| BEB | Bengali from Bangladesh |
| CDX | Chinese Dai in Xishuangbanna, China |
| CEU | Utah Residents (CEPH) with Northern and Western European Ancestry |
| CHB | Han Chinese in Beijing, China |
| CHS | Southern Han Chinese |
| CLM | Colombians from Medellin, Colombia |
| EAS | Eastern Asian |
| ESN | Esan in Nigeria |
| EUR | European |
| FIN | Finnish in Finland |
| GBR | British in England and Scotland |
| GIH | Gujarati Indian from Houston, Texas |
| GWD | Gambian in Western Divisions in the Gambia |
| IBS | Iberian Population in Spain |
| ITU | Indian Telugu from the UK |
| JPT | Japanese in Tokyo, Japan |
| KHV | Kinh in Ho Chi Minh City, Vietnam |
| LWK | Luhya in Webuye, Kenya |
| MSL | Mende in Sierra Leone |
| MXL | Mexican Ancestry from Los Angeles USA |
| NAT | Native American |
| OCE | Oceanian |
| PEL | Peruvians from Lima, Peru |
| PJL | Punjabi from Lahore, Pakistan |
| PUR | Puerto Ricans from Puerto Rico |
| SAS | Central/South Asian |
| STU | Sri Lankan Tamil from the UK |
| TSI | Toscani in Italia |
| WAS | Middle Eastern/Western Asian |
| YRI | Yoruba in Ibadan, Nigeria |

Table S1: Population code and description in the 1000 Genomes Project (TGP) and the Human Genome Diversity Project (HGDP) data.

| **Method** | **Parameters** |
|---|---|
| FLARE | min-mac=0 min-maf=0 ref=reference.vcf gt=query.vcf map=genetic_map.txt ref-panel=reference_population_label.txt out=output_basename |
| G-Nomix | query.vcf output_folder 18 False genetic_map.txt reference.vcf reference_population_label.txt default_config.yaml |
| Loter | -r reference_population_1.vcf reference_population_2.vcf reference_population_3.vcf -a query.vcf -f vcf -o output_result.txt -n 1 -v |
| MOSAIC | ADMIX Data/ -a 3 -n 200 -c 18 -p "AFR EAS EUR" -m 1 –nophase FALSE –singlePI TRUE |
| Recomb-Mix | -p reference.vcf -q query.vcf -a reference_population_label.txt -g genetic_map.txt -o output_folder -e 1.5 |
| RFMix | -f query.vcf -r reference.vcf -m reference_population_label.txt -g genetic_map.txt -o output_basename –chromosome=18 |
| SALAI-Net | –model-cp models/main_model/models/best_model.pth -q query.vcf -r reference.vcf -m reference_population_label.txt -o output_folder |

Table S2: Parameters of LAI methods used for performance analysis of the experiments. For Loter, it does not take sample map file "reference_population_label.txt" and reference file "reference.vcf". The reference file needs to be split into multiple reference file(s) per population as the input (e.g. reference_population_1.vcf, reference_population_2.vcf, etc.). For MOSAIC, the input files need to be converted to snpfile.CHR, rates.CHR, sample.names, and POPgenofile.CHR for each admixed and reference VCF files. All the files are located in the "Data" folder.

| **Method** | **100** | **250** | **500** | **1,000** |
|---|---|---|---|---|
| FLARE | 0.8664 | 0.9559 | 0.9894 | 0.9944 |
| G-Nomix | 0.9681 | 0.9882 | 0.9979 | 0.9989 |
| Loter | 0.8389 | 0.9482 | 0.9817 | 0.9909 |
| Recomb-Mix | 0.9919 | 0.9972 | 0.9995 | 0.9989 |
| RFMix | 0.8046 | 0.9733 | 0.9964 | 0.9982 |
| SALAI-Net | 0.9480 | 0.9936 | 0.9970 | 0.9970 |

Table S3: The squared Pearson's correlation coefficient $r^2$ with the reference panel sizes 100, 250, 500, and 1,000 of the three-way 15-generation inter-continental simulated datasets on FLARE, G-Nomix, Loter, Recomb-Mix, RFMix, and SALAI-Net. Markers were filtered with minor allele frequency $\leq 0.005$ and minor allele count $\leq 50$.

| **Method** | **15** | **50** | **100** | **200** |
|---|---|---|---|---|
| FLARE | 0.9894 | 0.9551 | 0.9159 | 0.8499 |
| G-Nomix | 0.9979 | 0.9813 | 0.9492 | 0.9204 |
| Loter | 0.9817 | 0.9461 | 0.8790 | 0.8130 |
| Recomb-Mix | 0.9995 | 0.9912 | 0.9725 | 0.9296 |
| RFMix | 0.9964 | 0.9451 | 0.7807 | 0.7520 |
| SALAI-Net | 0.9970 | 0.9762 | 0.8495 | 0.6435 |

Table S4: The squared Pearson's correlation coefficient $r^2$ with the generations 15, 50, 100, and 200 of the three-way 500-reference inter-continental simulated datasets on FLARE, G-Nomix, Loter, Recomb-Mix, RFMix, and SALAI-Net. Markers were filtered with minor allele frequency $\leq 0.005$ and minor allele count $\leq 50$.

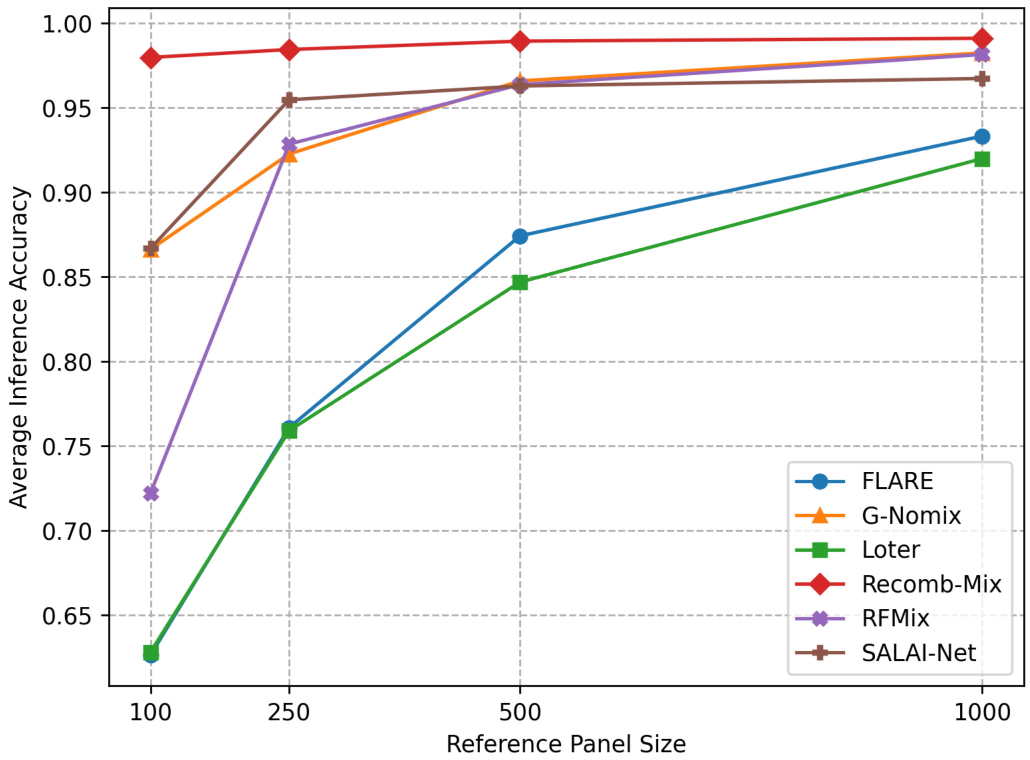


Figure S1: The average accuracy rates with the reference panel sizes 100, 250, 500, and 1,000 of the three-way 15-generation inter-continental simulated datasets on FLARE, G-Nomix, Loter, Recomb-Mix, RFMix, and SALAI-Net.

| **Method** | **100** | **250** | **500** | **1,000** |
|---|---|---|---|---|
| FLARE | 62.65 | 76.09 | 87.42 | 98.14 |
| G-Nomix | 86.63 | 92.26 | 96.58 | 98.24 |
| Loter | 62.82 | 75.91 | 84.69 | 91.99 |
| Recomb-Mix | 97.96 | 98.44 | 98.93 | 99.10 |
| RFMix | 72.22 | 92.85 | 96.36 | 98.14 |
| SALAI-Net | 86.69 | 95.47 | 96.28 | 96.72 |

Table S5: The average accuracy rates with the reference panel sizes 100, 250, 500, and 1,000 of the three-way 15-generation inter-continental simulated datasets on FLARE, G-Nomix, Loter, Recomb-Mix, RFMix, and SALAI-Net.

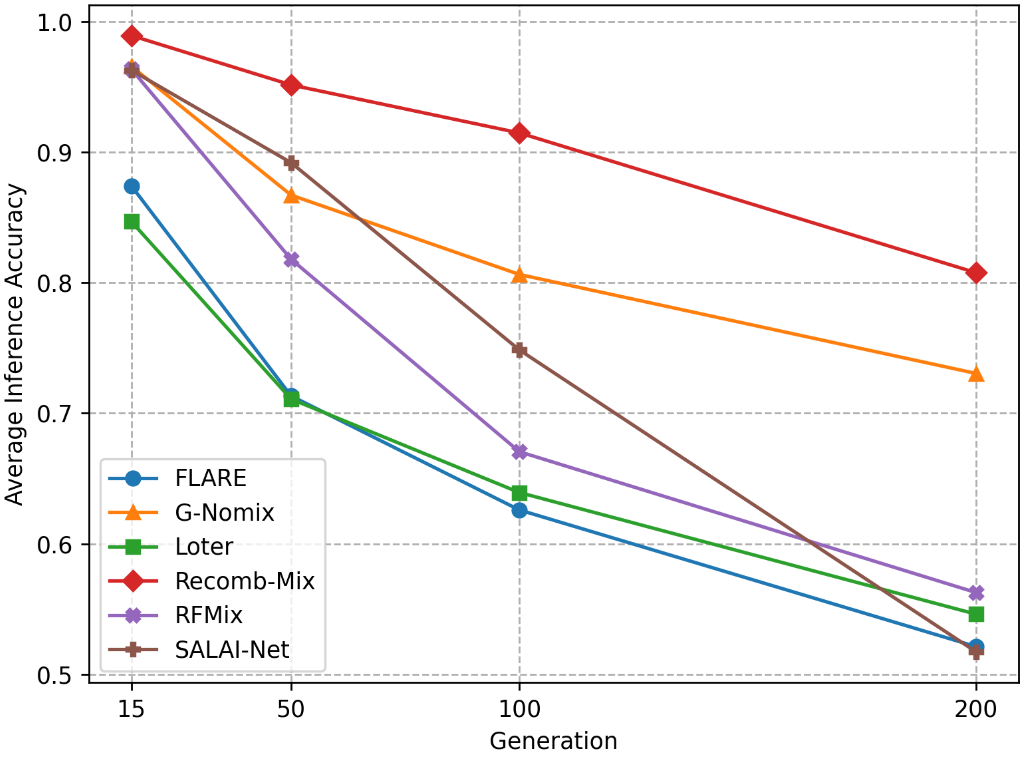


Figure S2: The average accuracy rates with the generations 15, 50, 100, and 200 of the three-way 500-reference inter-continental simulated datasets on FLARE, G-Nomix, Loter, Recomb-Mix, RFMix, and SALAI-Net.

| **Method** | **15** | **50** | **100** | **200** |
|---|---|---|---|---|
| FLARE | 87.42 | 71.32 | 62.59 | 52.12 |
| G-Nomix | 96.58 | 86.71 | 80.63 | 73.04 |
| Loter | 84.69 | 71.08 | 63.92 | 54.60 |
| Recomb-Mix | 98.93 | 95.15 | 91.47 | 80.76 |
| RFMix | 96.36 | 81.79 | 67.05 | 56.26 |
| SALAI-Net | 96.28 | 89.19 | 74.84 | 51.72 |

Table S6: The average accuracy rates with the generations 15, 50, 100, and 200 of the three-way 500-reference inter-continental simulated datasets on FLARE, G-Nomix, Loter, Recomb-Mix, RFMix, and SALAI-Net.

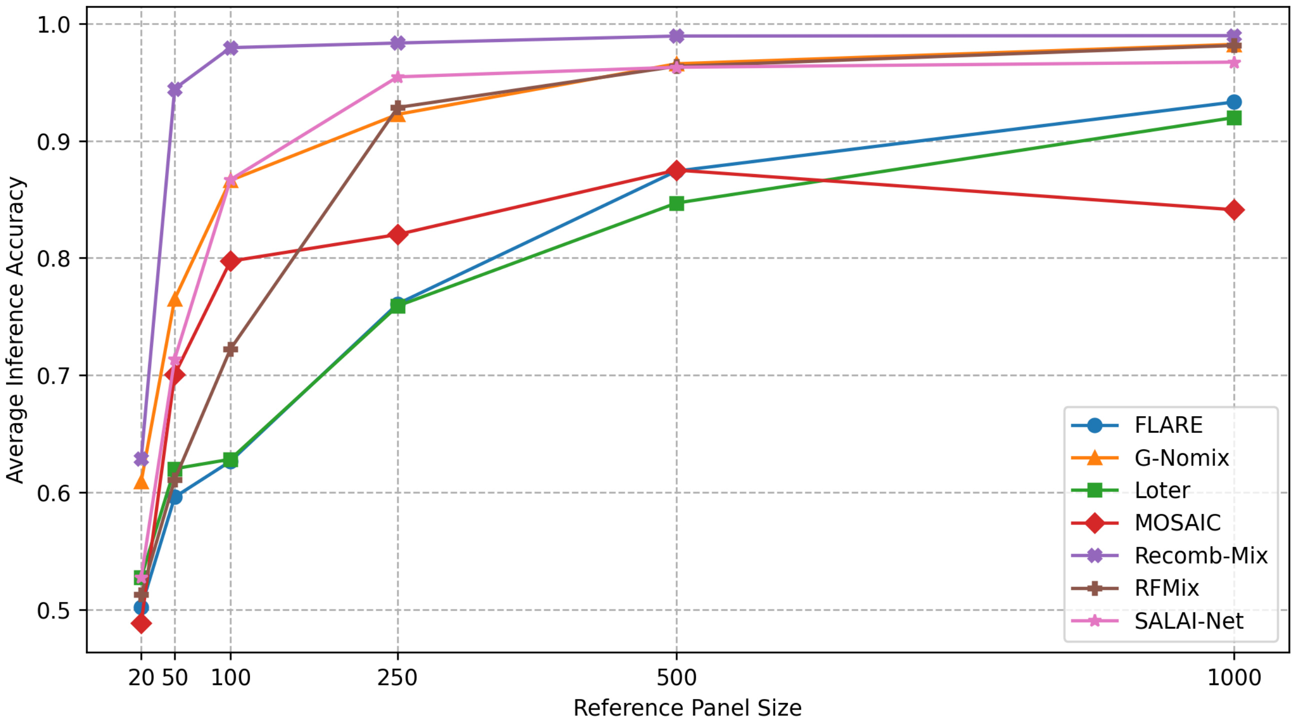


Figure S3: The average accuracy rates with the reference panel sizes 20, 50, 100, 250, 500, and 1,000 of the three-way 15-generation inter-continental simulated datasets on FLARE, G-Nomix, Loter, MOSAIC, Recomb-Mix, RFMix, and SALAI-Net.

| **Method** | **20** | **50** | **100** | **250** | **500** | **1,000** |
|---|---|---|---|---|---|---|
| FLARE | 50.21 | 59.59 | 62.65 | 76.09 | 87.42 | 93.32 |
| G-Nomix | 60.92 | 76.50 | 86.63 | 92.26 | 96.58 | 98.24 |
| Loter | 52.75 | 62.02 | 62.82 | 75.91 | 84.69 | 91.99 |
| MOSAIC | 48.84 | 70.06 | 79.73 | 82.01 | 87.51 | 84.12 |
| Recomb-Mix | 62.85 | 94.45 | 97.96 | 98.35 | 98.95 | 98.98 |
| RFMix | 51.28 | 61.08 | 72.22 | 92.85 | 96.36 | 98.14 |
| SALAI-Net | 52.72 | 71.39 | 86.69 | 95.47 | 96.28 | 96.72 |

Table S7: The average accuracy rates with the reference panel sizes 20, 50, 100, 250, 500, and 1,000 of the three-way 15-generation inter-continental simulated datasets on FLARE, G-Nomix, Loter, MOSAIC, Recomb-Mix, RFMix, and SALAI-Net.

| **Method** | **250** | **500** | **1,000** |
|---|---|---|---|
| FLARE | 0.6484 | 0.8509 | 0.8788 |
| G-Nomix | 0.7563 | 0.9100 | 0.9812 |
| Loter | 0.6457 | 0.7705 | 0.8240 |
| Recomb-Mix | 0.9544 | 0.9798 | 0.9904 |
| RFMix | 0.5107 | 0.8699 | 0.9002 |
| SALAI-Net | 0.7407 | 0.9535 | 0.9740 |

Table S8: The squared Pearson's correlation coefficient $r^2$ with the reference panel sizes 250, 500, and 1,000 of the seven-way 15-generation inter-continental simulated datasets on FLARE, G-Nomix, Loter, Recomb-Mix, RFMix, and SALAI-Net. Markers were filtered with minor allele frequency $\leq 0.005$ and minor allele count $\leq 50$. The reference panel size 100 case was not included because the number of markers was too small and may have influenced the outcome after the filtering.

| **Method** | **15** | **50** | **100** | **200** |
|---|---|---|---|---|
| FLARE | 0.8509 | 0.8323 | 0.7831 | 0.6558 |
| G-Nomix | 0.9100 | 0.8322 | 0.8462 | 0.5845 |
| Loter | 0.7705 | 0.7694 | 0.6970 | 0.5570 |
| Recomb-Mix | 0.9798 | 0.9268 | 0.8728 | 0.5553 |
| RFMix | 0.8699 | 0.6028 | 0.5968 | 0.4187 |
| SALAI-Net | 0.9535 | 0.7469 | 0.5457 | 0.4869 |

Table S9: The squared Pearson's correlation coefficient $r^2$ with the generations 15, 50, 100, and 200 of the seven-way 500-reference inter-continental simulated datasets on FLARE, G-Nomix, Loter, Recomb-Mix, RFMix, and SALAI-Net. Markers were filtered with minor allele frequency $\leq 0.005$ and minor allele count $\leq 50$.

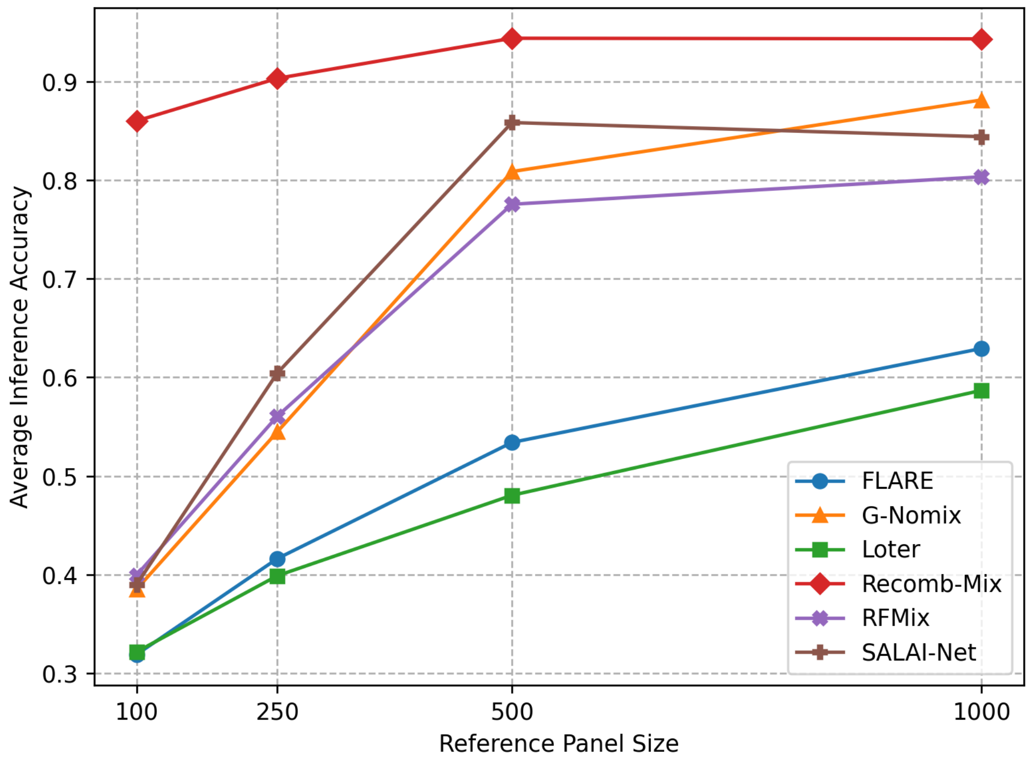


Figure S4: The average accuracy rates with the reference panel sizes 100, 250, 500, and 1,000 of the seven-way 15-generation inter-continental simulated datasets on FLARE, G-Nomix, Loter, Recomb-Mix, RFMix, and SALAI-Net.

| **Method** | **100** | **250** | **500** | **1,000** |
|---|---|---|---|---|
| FLARE | 31.89 | 41.62 | 53.41 | 62.93 |
| G-Nomix | 38.50 | 54.48 | 80.88 | 88.14 |
| Loter | 32.16 | 39.88 | 48.05 | 58.68 |
| Recomb-Mix | 86.02 | 90.33 | 94.40 | 94.34 |
| RFMix | 39.93 | 56.06 | 77.56 | 80.35 |
| SALAI-Net | 38.99 | 60.44 | 85.85 | 84.42 |

Table S10: The average accuracy rates with the reference panel sizes 100, 250, 500, and 1,000 of the seven-way 15-generation inter-continental simulated datasets on FLARE, G-Nomix, Loter, Recomb-Mix, RFMix, and SALAI-Net.

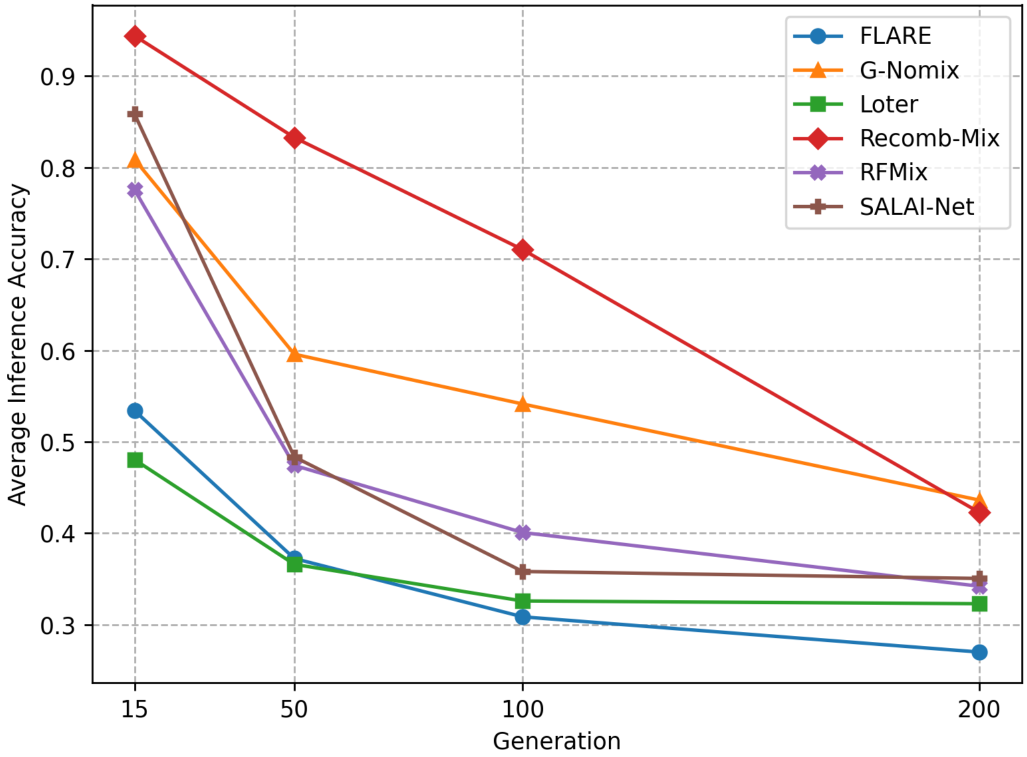


Figure S5: The average accuracy rates with the generations 15, 50, 100, and 200 of the seven-way 500-reference inter-continental simulated datasets on FLARE, G-Nomix, Loter, Recomb-Mix, RFMix, and SALAI-Net.

| **Method** | **15** | **50** | **100** | **200** |
|---|---|---|---|---|
| FLARE | 53.41 | 37.23 | 30.85 | 27.00 |
| G-Nomix | 80.88 | 59.58 | 54.13 | 43.62 |
| Loter | 48.05 | 36.59 | 32.58 | 32.29 |
| Recomb-Mix | 94.40 | 83.27 | 71.01 | 42.27 |
| RFMix | 77.56 | 47.41 | 40.07 | 34.20 |
| SALAI-Net | 85.85 | 48.34 | 35.81 | 35.05 |

Table S11: The average accuracy rates with the generations 15, 50, 100, and 200 of the seven-way 500-reference inter-continental simulated datasets on FLARE, G-Nomix, Loter, Recomb-Mix, RFMix, and SALAI-Net.

| **Method** | **250** | **500** | **1,000** |
|---|---|---|---|
| FLARE | 0.7232 | 0.7538 | 0.9273 |
| G-Nomix | 0.8560 | 0.9235 | 0.9820 |
| Loter | 0.6441 | 0.7264 | 0.8794 |
| Recomb-Mix | 0.9299 | 0.9625 | 0.9800 |
| RFMix | 0.7180 | 0.8024 | 0.9077 |
| SALAI-Net | 0.7910 | 0.9081 | 0.9687 |

Table S12: The squared Pearson's correlation coefficient $r^2$ with the reference panel sizes 250, 500, and 1,000 of the three-way 15-generation intra-continental simulated datasets on FLARE, G-Nomix, Loter, Recomb-Mix, RFMix, and SALAI-Net. Markers were filtered with minor allele frequency $\leq 0.005$ and minor allele count $\leq 50$. The reference panel size 100 case was not included because the number of markers was too small and may have influenced the outcome after the filtering.

| **Method** | **15** | **50** | **100** | **200** |
|---|---|---|---|---|
| FLARE | 0.7538 | 0.7768 | 0.7784 | 0.7595 |
| G-Nomix | 0.9235 | 0.7912 | 0.7992 | 0.8048 |
| Loter | 0.7264 | 0.6406 | 0.6978 | 0.6993 |
| Recomb-Mix | 0.9625 | 0.8506 | 0.7930 | 0.7428 |
| RFMix | 0.8024 | 0.7072 | 0.6686 | 0.6035 |
| SALAI-Net | 0.9081 | 0.7281 | 0.6269 | 0.4679 |

Table S13: The squared Pearson's correlation coefficient $r^2$ with the generations 15, 50, 100, and 200 of the three-way 500-reference intra-continental simulated datasets on FLARE, G-Nomix, Loter, Recomb-Mix, RFMix, and SALAI-Net. Markers were filtered with minor allele frequency $\leq 0.005$ and minor allele count $\leq 50$.

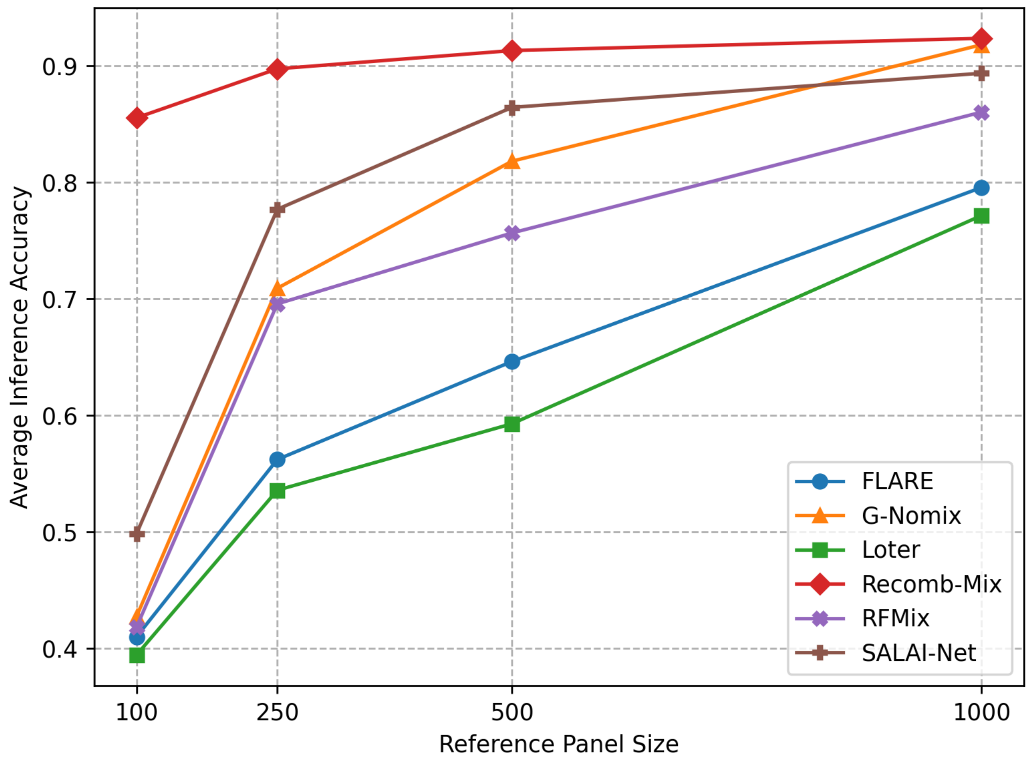


Figure S6: The average accuracy rates with the reference panel sizes 100, 250, 500, and 1,000 of the three-way 15-generation intra-continental simulated datasets on FLARE, G-Nomix, Loter, Recomb-Mix, RFMix, and SALAI-Net.

| **Method** | **100** | **250** | **500** | **1,000** |
|---|---|---|---|---|
| FLARE | 40.98 | 56.21 | 64.62 | 79.56 |
| G-Nomix | 42.71 | 70.91 | 81.83 | 91.82 |
| Loter | 39.42 | 53.57 | 59.28 | 77.15 |
| Recomb-Mix | 85.55 | 89.76 | 91.33 | 92.38 |
| RFMix | 41.81 | 69.57 | 75.66 | 86.04 |
| SALAI-Net | 49.83 | 77.70 | 86.45 | 89.38 |

Table S14: The average accuracy rates with the reference panel sizes 100, 250, 500, and 1,000 of the three-way 15-generation intra-continental simulated datasets on FLARE, G-Nomix, Loter, Recomb-Mix, RFMix, and SALAI-Net.

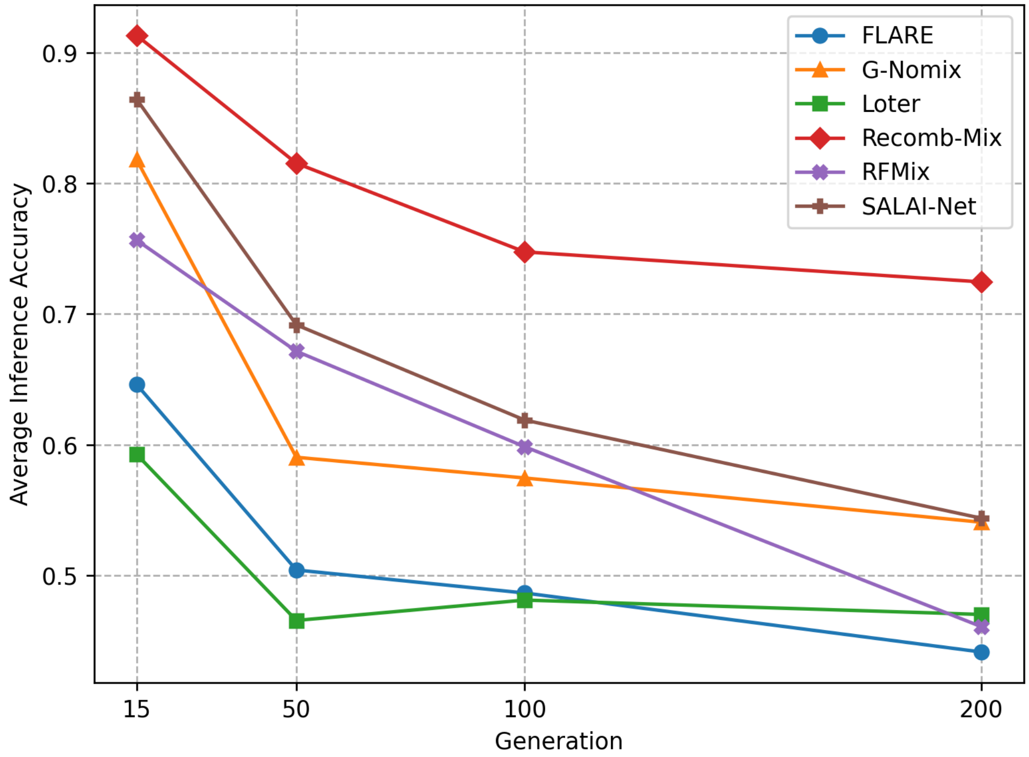


Figure S7: The average accuracy rates with the generations 15, 50, 100, and 200 of the three-way 500-reference intra-continental simulated datasets on FLARE, G-Nomix, Loter, Recomb-Mix, RFMix, and SALAI-Net.

| **Method** | **15** | **50** | **100** | **200** |
|---|---|---|---|---|
| FLARE | 64.62 | 50.42 | 48.66 | 44.15 |
| G-Nomix | 81.83 | 59.04 | 57.46 | 54.07 |
| Loter | 59.28 | 46.56 | 48.12 | 47.02 |
| Recomb-Mix | 91.33 | 81.54 | 74.76 | 72.46 |
| RFMix | 75.66 | 67.16 | 59.85 | 46.09 |
| SALAI-Net | 86.45 | 69.18 | 61.89 | 54.38 |

Table S15: The average accuracy rates with the generations 15, 50, 100, and 200 of the three-way 500-reference intra-continental simulated datasets on FLARE, G-Nomix, Loter, Recomb-Mix, RFMix, and SALAI-Net.

| Method | Even Founders and References | Uneven Founders | Uneven References |
|---|---|---|---|
| Loter | 59.28 | 51.67 | 52.98 |
| FLARE | 64.62 | 59.60 | 56.85 |
| RFMix | 75.66 | 74.61 | 72.47 |
| G-Nomix | 81.83 | 80.16 | 73.88 |
| SALAI-Net | 86.45 | 83.07 | 79.49 |
| Recomb-Mix | **91.33** | **89.20** | **83.61** |

Table S16: The average accuracy rates of FLARE, G-Nomix, Loter, Recomb-Mix, RFMix, and SALAI-Net performing LAI on three-way 15-generation 500-reference intra-continental simulated datasets with even or uneven number of individuals per population in founder or reference panel.

| Method | 15 | 50 | 100 | 200 |
|---|---|---|---|---|
| FLARE | 0.8811 | 0.9019 | 0.8764 | 0.8731 |
| G-Nomix | 0.8799 | 0.9012 | 0.8746 | 0.8278 |
| Loter | 0.8570 | 0.8525 | 0.8653 | 0.7784 |
| Recomb-Mix | 0.8827 | 0.9188 | 0.8981 | 0.7762 |
| RFMix | 0.9556 | 0.8195 | 0.6965 | 0.6590 |
| SALAI-Net | 0.8923 | 0.8504 | 0.7447 | 0.5433 |

Table S17: The squared Pearson's correlation coefficient $r^2$ with the generations 15, 50, 100, and 200 of the three-way 500-misspecified-reference inter-continental simulated datasets on FLARE, G-Nomix, Loter, Recomb-Mix, RFMix, and SALAI-Net. Markers were filtered with minor allele frequency $\leq 0.005$ and minor allele count $\leq 50$.

| Method | 15 | 50 | 100 | 200 |
|---|---|---|---|---|
| FLARE | 66.16 | 54.29 | 47.44 | 42.65 |
| G-Nomix | 82.46 | 74.42 | 69.45 | 63.10 |
| Loter | 62.71 | 55.11 | 50.08 | 44.71 |
| Recomb-Mix | 85.15 | 86.93 | 80.99 | 66.08 |
| RFMix | 91.36 | 69.15 | 53.82 | 49.21 |
| SALAI-Net | 81.00 | 74.26 | 61.62 | 49.73 |

Table S18: The average accuracy rates with the generations 15, 50, 100, and 200 of the three-way 500-misspecified-reference inter-continental simulated datasets on FLARE, G-Nomix, Loter, Recomb-Mix, RFMix, and SALAI-Net.

| Method | No Phasing Error | Phasing Error on Targets | Phasing Error on References |
|---|---|---|---|
| Loter | 64.00 | 62.72 | 63.35 |
| FLARE | 63.68 | 63.61 | 64.15 |
| RFMix | 72.52 | 71.26 | 72.77 |
| SALAI-Net | 86.81 | 85.82 | 87.54 |
| G-Nomix | 86.70 | 86.37 | 86.54 |
| Recomb-Mix | 97.97 | 97.39 | 97.98 |

Table S19: The diploid accuracy rates of FLARE, G-Nomix, Loter, Recomb-Mix, RFMix, and SALAI-Net performing LAI on three-way 15-generation 100-reference inter-continental simulated datasets with no phasing error, phasing error on target panel, and phasing error on reference panel.

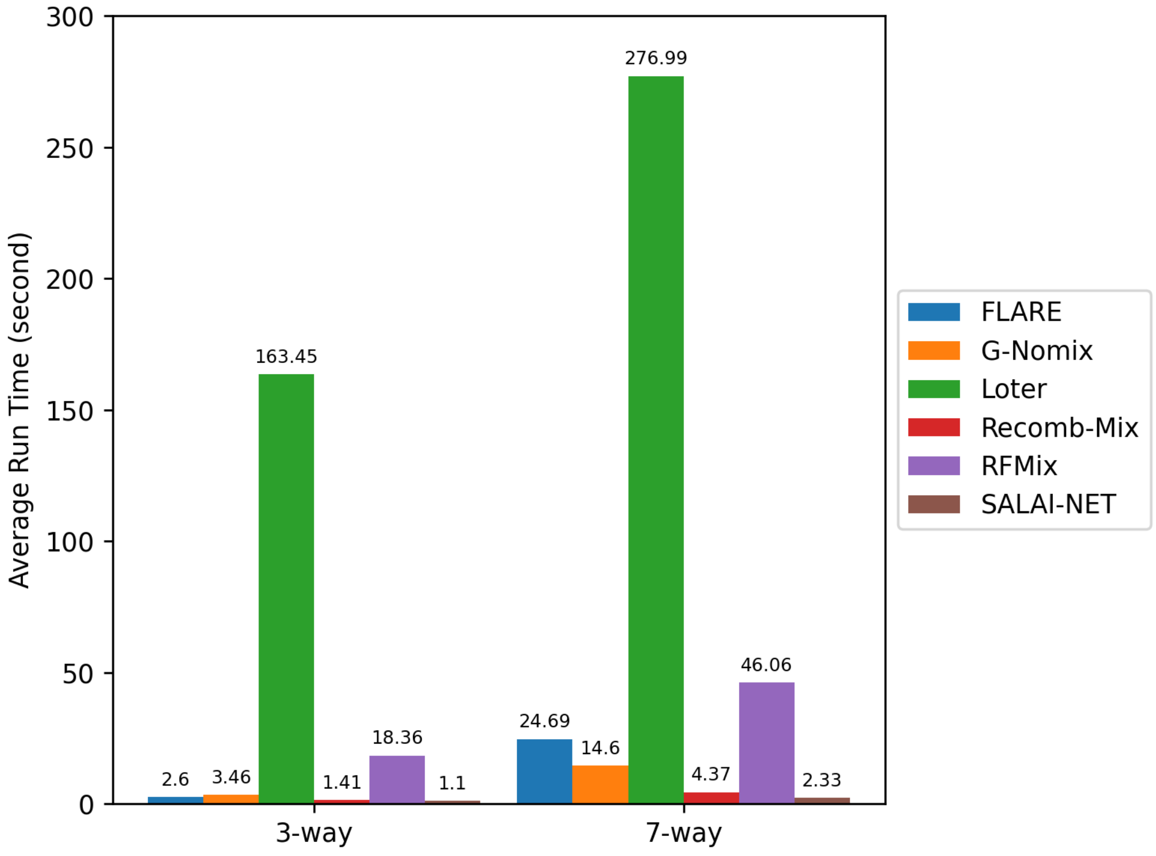


Figure S8: The average run time (second) of LAI methods FLARE, G-Nomix, Loter, Recomb-Mix, RFMix, and SALAI-Net for querying local ancestry information of an admixed individual haplotype on three-way and seven-way reference panels.

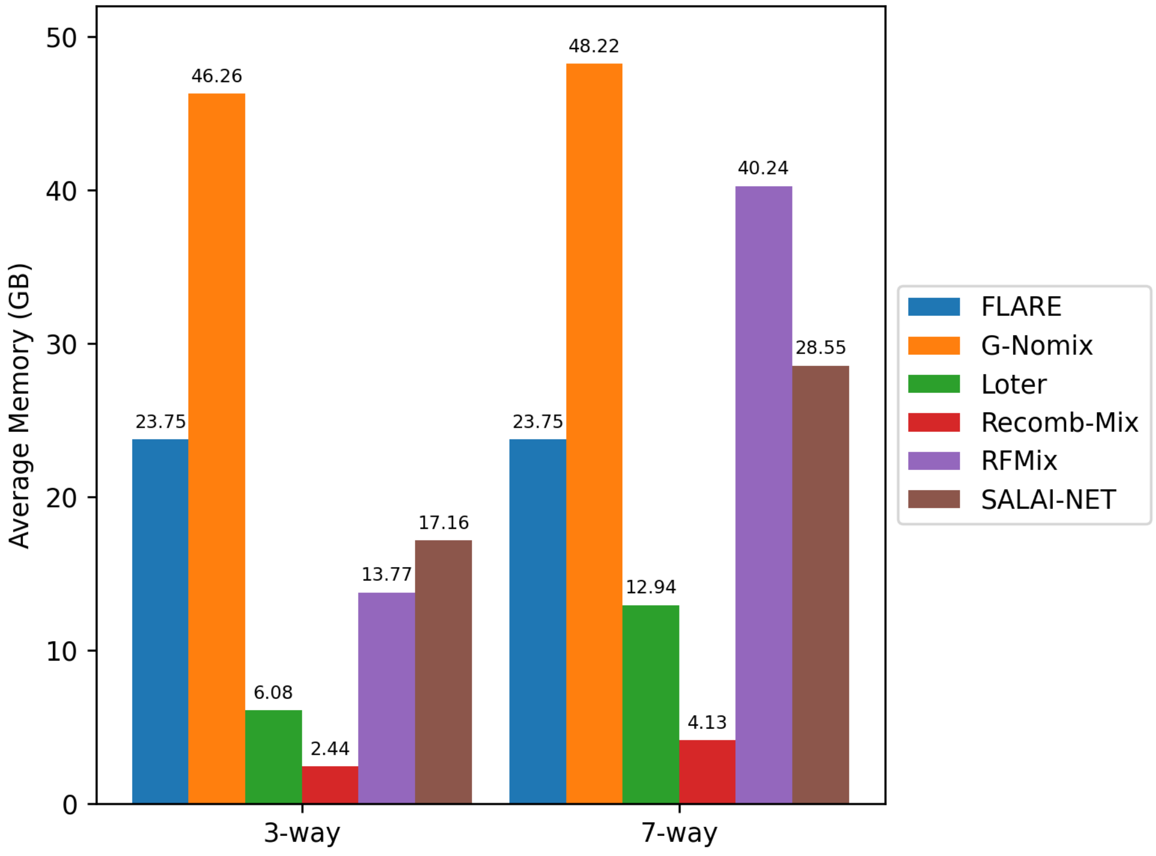


Figure S9: The average memory (GB) of LAI methods FLARE, G-Nomix, Loter, Recomb-Mix, RFMix, and SALAI-Net for querying local ancestry information of an admixed individual haplotype on three-way and seven-way reference panels.

| **Method** | **3-way Run Time (second)** | **7-way Run Time (second)** | **3-way Memory (GB)** | **7-way Memory (GB)** |
|---|---|---|---|---|
| FLARE | 2.60 | 24.69 | 23.75 | 23.75 |
| G-Nomix | 3.46 | 14.60 | 46.26 | 48.22 |
| Loter | 163.45 | 276.99 | 6.08 | 12.94 |
| Recomb-Mix | 1.41 | 4.37 | 2.44 | 4.13 |
| RFMix | 18.36 | 46.06 | 13.77 | 40.24 |
| SALAI-Net | 1.10 | 2.33 | 17.16 | 28.55 |

Table S20: The average run time (second) and maximum amount of physical memory (GB) of LAI methods FLARE, G-Nomix, Loter, Recomb-Mix, RFMix, and SALAI-Net for querying local ancestry information of an admixed individual haplotype. Values were averaged from runs of inter- and intra-continental, with reference panel sizes 100, 250, 500, and 1,000 and generations 15, 50, 100, and 200. All methods were tested on a single-node machine with an Intel Xeon Gold 5215 2.50 GHz processor and 200 gigabytes of RAM.

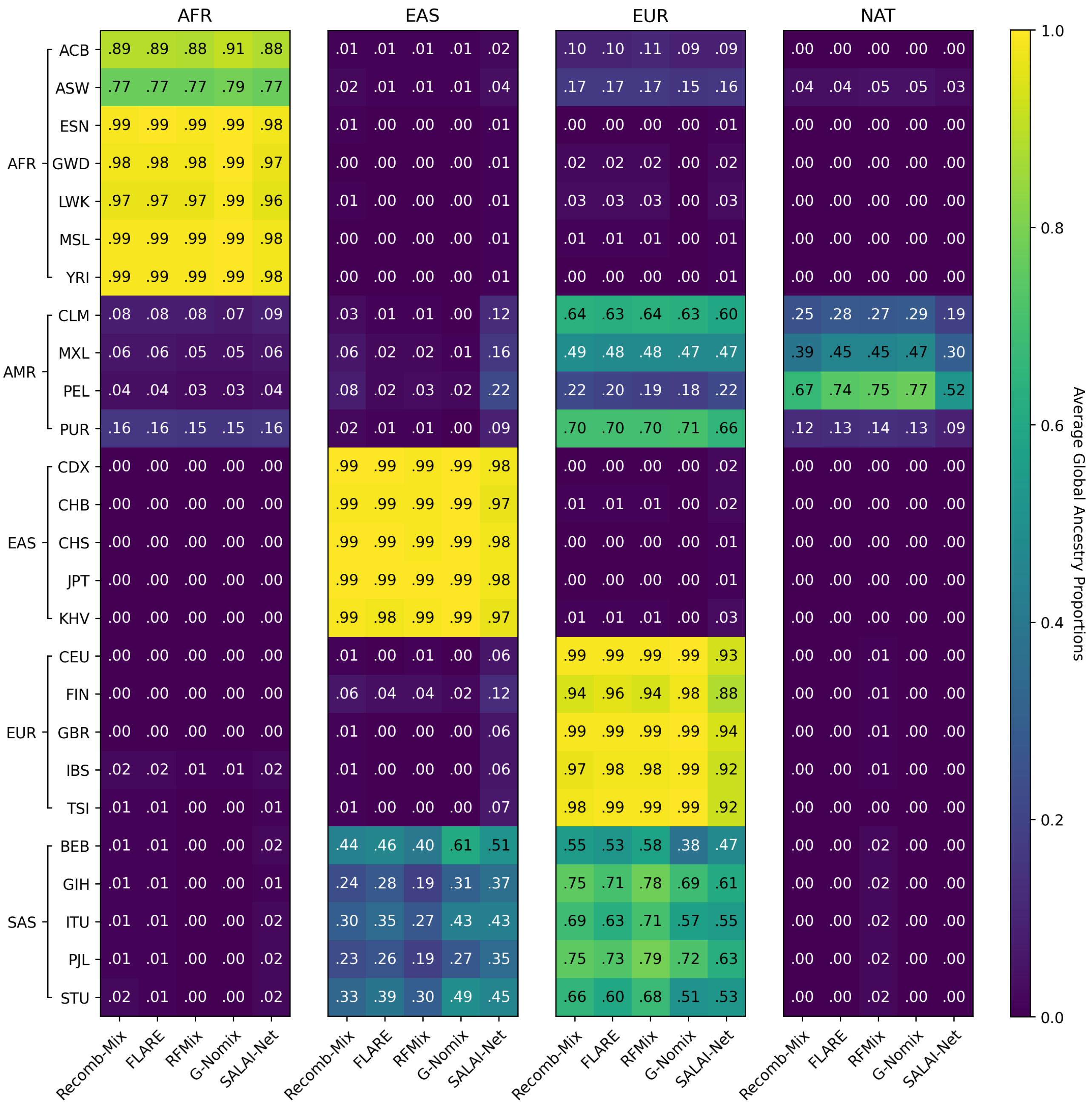


Figure S10: The average global ancestry proportions in the 1000 Genomes Project (TGP) Chromosome 18 data using four reference ancestries from the Human Genome Diversity Project (HGDP) data. The methods are Recomb-Mix, FLARE, RFMix, G-Nomix, and SALAI-Net. Descriptions of the populations are in Supplemental Table S1.

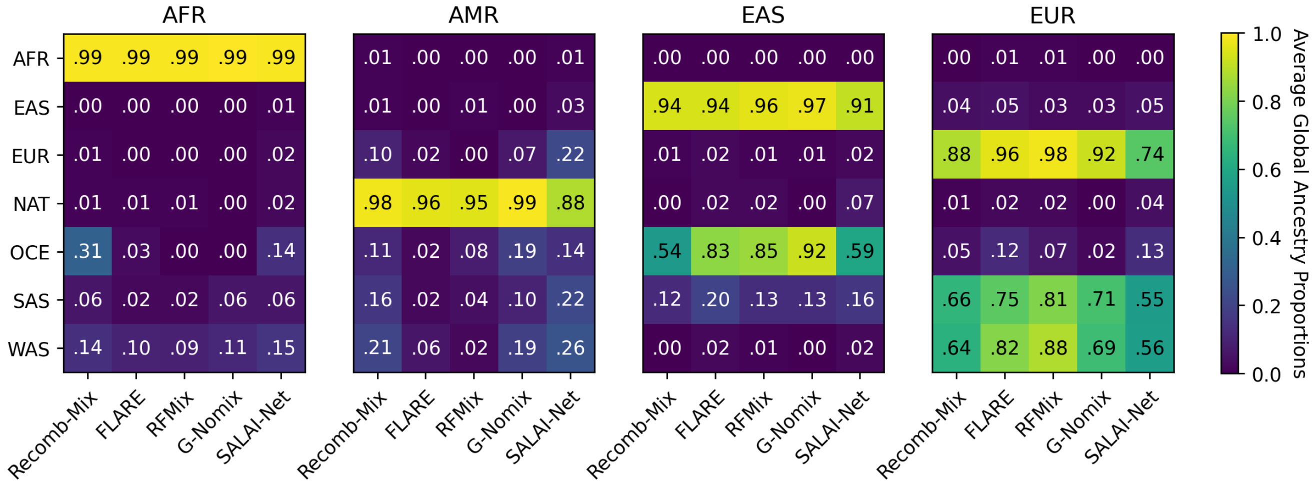


Figure S11: The average global ancestry proportions in the Human Genome Diversity Project (HGDP) Chromosome 18 data using four reference ancestries from the 1000 Genomes Project (TGP) data. The methods are Recomb-Mix, FLARE, RFMix, G-Nomix, and SALAI-Net. Descriptions of the populations are in Supplemental Table S1.

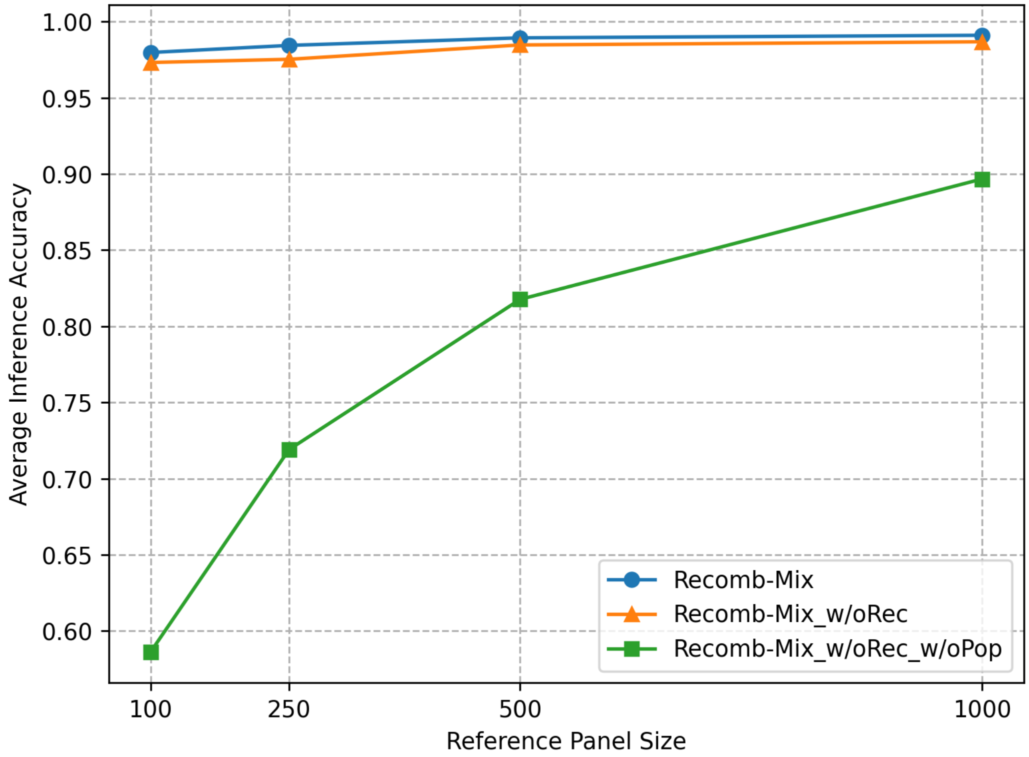


Figure S12: The average accuracy rates with reference panel sizes 100, 250, 500, and 1,000 of the three-way 15-generation inter-continental simulated datasets. Recomb-Mix_w/oRec method is Recomb-Mix without using recombination rates in the objective function. Instead, a constant value is used as the template change penalty. Recomb-Mix_w/oRec_w/oPop is Recomb-Mix without using recombination rates in the objective function and without considering the zero template change penalty within each population. Instead, the template change penalty occurs when the threading path is changed to a different haplotype template from the current one, regardless of their population labels, as Loter did.

| **Method** | **100** | **250** | **500** | **1,000** |
|---|---|---|---|---|
| Recomb-Mix | 97.97 | 98.44 | 98.93 | 99.10 |
| Recomb-Mix_w/oRec | 97.32 | 97.53 | 98.47 | 98.68 |
| Recomb-Mix_w/oRec_w/oPop | 58.62 | 71.91 | 81.77 | 89.67 |

Table S21: The average accuracy rates of LAI methods Recomb-Mix, Recomb-Mix_w/oRec, and Recomb-Mix_w/oRec_w/oPop on the three-way 15-generation inter-continental simulated datasets with reference panel sizes 100, 250, 500, and 1,000.

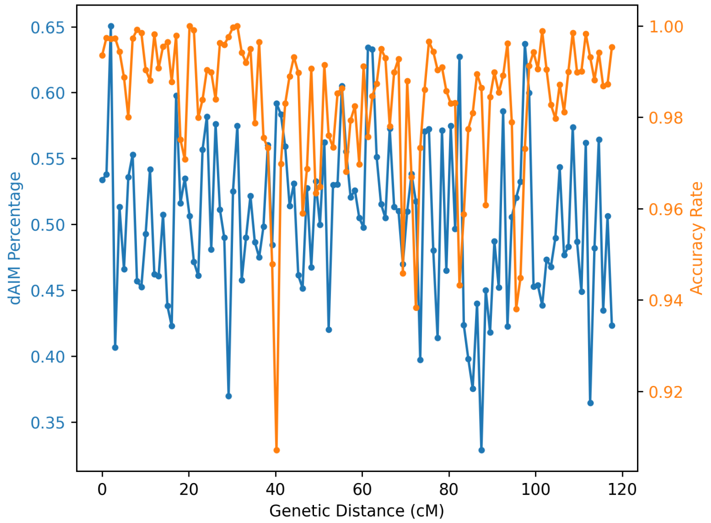


Figure S13: The discrete AIM (dAIM) density and local ancestry inference accuracy rate of a three-way 15-generation 100-reference inter-continental Chromosome 18 simulated dataset. Each bin is 1 centiMorgan (cM), showing the markers' dAIM percentage and the average accuracy rate of local ancestry inferred by Recomb-Mix.

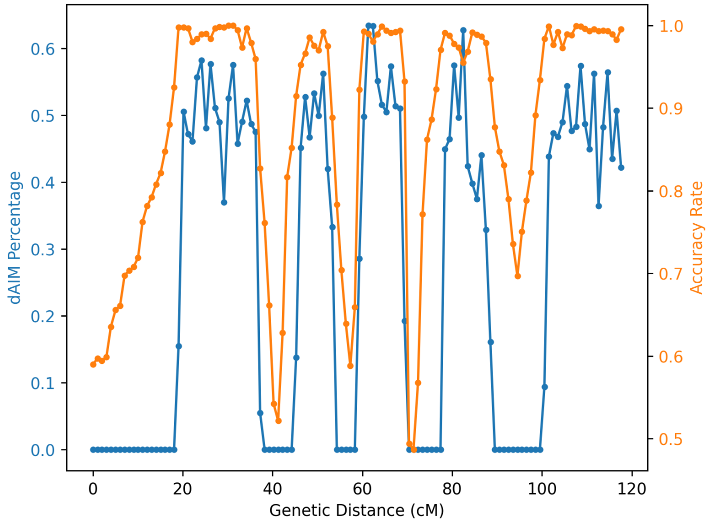


Figure S14: The discrete AIM (dAIM) density and local ancestry inference accuracy rate of a three-way 15-generation 100-reference inter-continental Chromosome 18 simulated dataset. The dataset was engineered to have certain regions of dAIMs removed. Each bin is 1 centiMorgan (cM), showing the markers' dAIM percentage and the average accuracy rate of local ancestry inferred by Recomb-Mix.

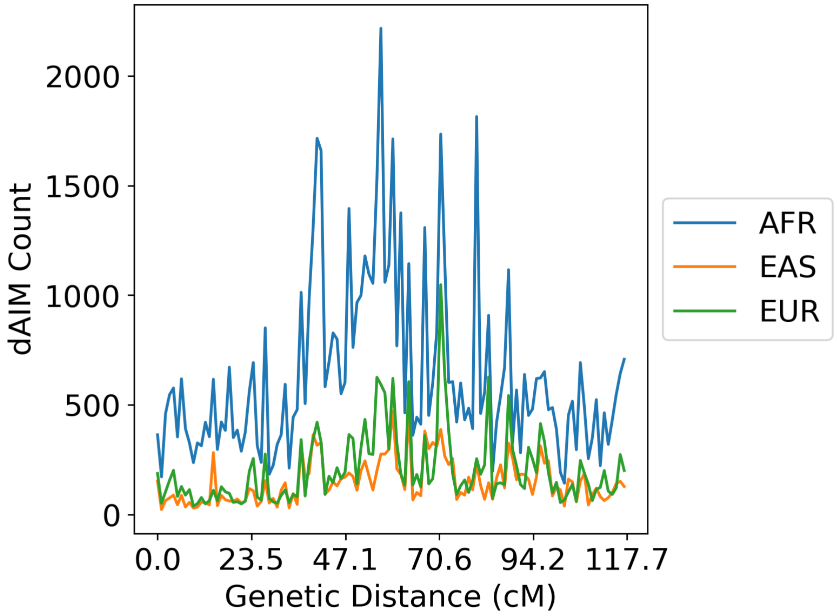


Figure S15: The number of dAIMs in a three-way 15-generation 100-reference inter-continental sequencing dataset.

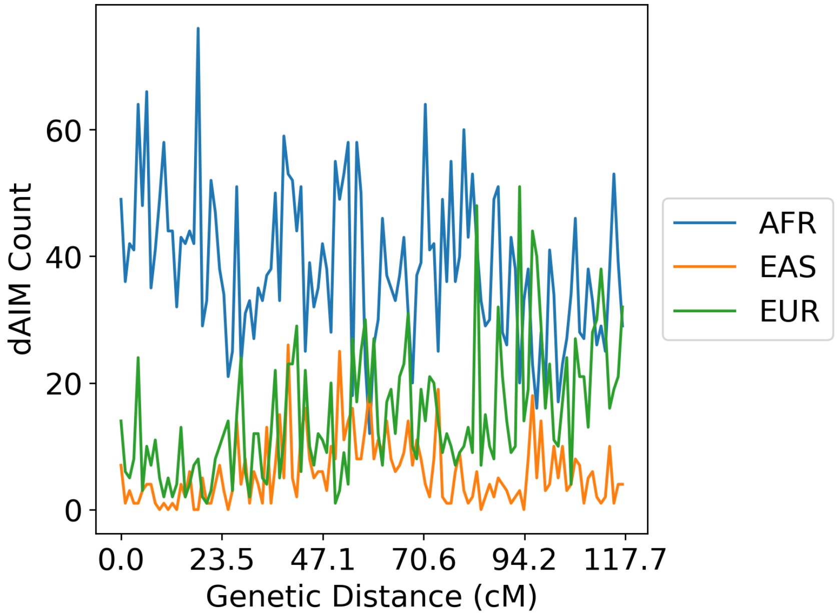


Figure S16: The number of dAIMs in a three-way 15-generation 100-reference inter-continental genotyping dataset.

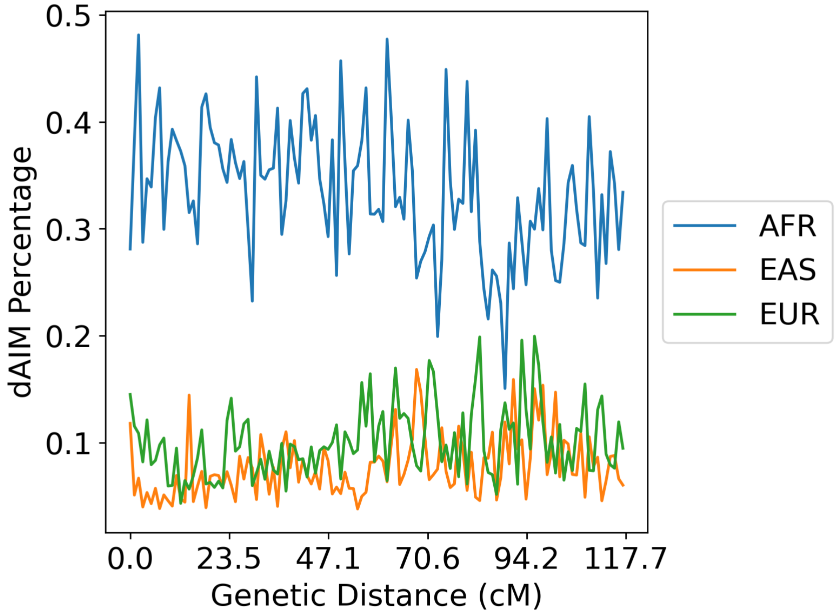


Figure S17: The dAIM density in a three-way 15-generation 100-reference inter-continental sequencing dataset.

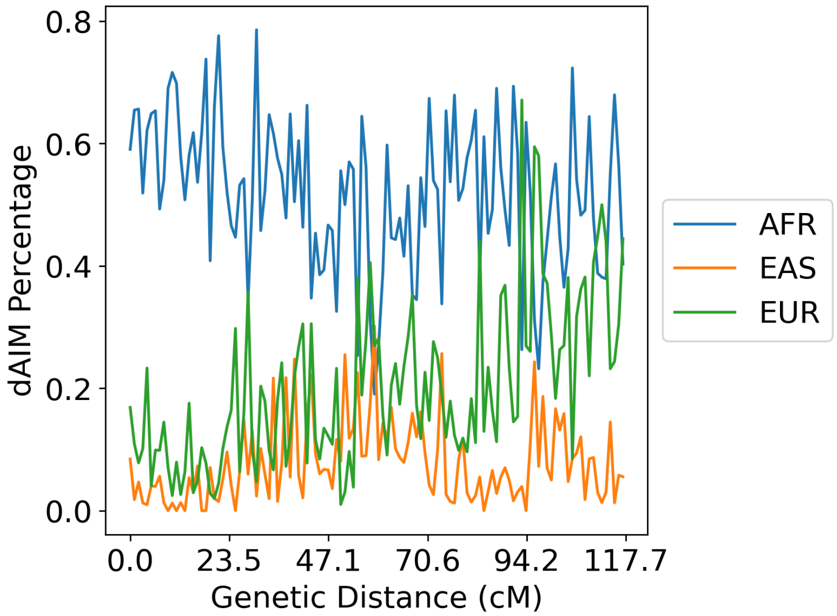


Figure S18: The dAIM density in a three-way 15-generation 100-reference inter-continental genotyping dataset.
